# EML4-ALK V3 drives cell migration through NEK9 and NEK7 kinases in non-small-cell lung cancer

**DOI:** 10.1101/567305

**Authors:** Laura O’Regan, Giancarlo Barone, Rozita Adib, Chang Gok Woo, Hui Jeong Jeong, Emily L. Richardson, Mark W. Richards, Patricia A.J. Muller, Spencer J. Collis, Dean A. Fennell, Jene Choi, Richard Bayliss, Andrew M. Fry

**Affiliations:** Department of Molecular and Cell Biology, University of Leicester, Lancaster Road, Leicester LE1 9HN, U.K.; Department of Oncology and Metabolism, Sheffield Institute for Nucleic Acids (SInFoNiA), University of Sheffield, Beech Hill Road, Sheffield S10 2RX, U.K.; Department of Pathology, Chungbuk National University Hospital, Chungbuk National University College of Medicine, Cheongju, Korea.; Department of Pathology, Asan Medical Center, University of Ulsan College of Medicine, Seoul, Korea.; School of Molecular and Cellular Biology, Astbury Centre for Structural Molecular Biology, Faculty of Biological Sciences, University of Leeds, Leeds LS2 9JT, U.K.; Cancer Research UK Manchester Institute, University of Manchester, Alderley Park SK10 4TG, U.K.; Cancer Research Centre, University of Leicester, Robert Kilpatrick Clinical Sciences Building, Leicester LE1 9HN, U.K.

**Keywords:** EML4-ALK, EML4, NEK9, NEK7, NSCLC, microtubules, cell migration, metastasis

## Abstract

EML4-ALK is an oncogenic fusion present in ∼5% lung adenocarcinomas. However, distinct EML4-ALK variants differ in the length of the EML4 microtubule-associated protein encoded within the fusion and are associated with a poorly understood variability in disease progression and therapeutic response. Here, we show that EML4-ALK variant 3, which is linked to accelerated metastatic spread and worse patient outcome, causes microtubule stabilization, formation of extended cytoplasmic protrusions, loss of cell polarity and increased cell migration. Strikingly, this is dependent upon the NEK9 kinase that interacts with the N-terminal region of EML4. Overexpression of wild-type EML4, as well as constitutive activation of NEK9, also perturbs cell morphology and accelerates cell migration in a manner that requires the downstream kinase NEK7 but not ALK activity. Moreover, elevated NEK9 is associated in patients with EML4-ALK V3 expression, as well as reduced progression-free and overall survival. Hence, we propose that EML4-ALK V3 promotes microtubule stabilization through recruitment of NEK9 and NEK7 to increase cell migration and that this represents a novel actionable pathway that drives disease progression in lung cancer.

## INTRODUCTION

The EML4-ALK translocation is an oncogenic driver in a subset of lung cancers that results from an inversion on the short arm of chromosome 2 fusing an N-terminal fragment of the echinoderm microtubule-associated protein-like 4, EML4^1, 2^, to the C-terminal tyrosine kinase domain of the anaplastic lymphoma kinase, ALK. The fusion was first identified in non-small-cell lung cancer (NSCLC), where it is present in approximately 5% of cases, but has since been identified in other tumour types, including breast and colorectal cancers^3, 4, 5^. The majority of EML4-ALK lung cancers, which are predominantly adenocarcinomas, respond remarkably well to catalytic inhibitors of the ALK tyrosine kinase, such as crizotinib. However, this approach is not curative as acquired resistance to ALK inhibitors is inevitable due to either secondary mutations in the ALK tyrosine kinase domain or off-target alterations that switch dependence to other signalling pathways^6,7,8,9,10,11^. Consequently, effective therapies capable of selectively targeting ALK inhibitor resistant lung cancers are warranted.

It is also now clear that not all EML4-ALK patients respond well to ALK inhibitor therapy in the first place^12^. One potential explanation for this is the presence of alternative EML4-ALK variants that arise from distinct breakpoints^13, 14^. All fusions encode the full C-terminal catalytic domain of the ALK kinase and an N-terminal oligomerization domain from the EML4 protein that promotes autophosphorylation. However, alternative breakpoints in the *EML4* gene lead to variable amounts of the EML4 protein being present in the fusions. The N-terminal oligomerization domain of EML4 has been shown by X-ray crystallography to form trimers^15^. This sequence is followed by an unstructured region of approximately 150 residues rich in serine, threonine and basic residues. Based on crystallographic analysis of the related EML1 protein, the ∼600 residue C-terminal region folds into a tandem pair of atypical β-propellers termed the TAPE domain^16^. Structure-function studies have shown that while the C-terminal TAPE domain binds to α*/*β-tubulin heterodimers, it is the N-terminal domain (NTD) encompassing the trimerization motif and unstructured region that promotes binding to polymerized microtubules^15, 16^.

While all EML4-ALK fusion proteins have the trimerization motif, the distinct breakpoints mean that the different variants encode more or less of the unstructured and TAPE domains. For example, the longer variants, 1 and 2, encode the unstructured domain and a partial fragment of the TAPE domain, while the shorter variants, 3a, 3b and 5, have none of the TAPE domain. Interestingly, the presence of a TAPE domain fragment renders the fusion protein unstable and dependent on the HSP90 chaperone for expression^16, 17^. Indeed, binding to HSP90 may interfere with microtubule association as these longer variants localise poorly to the microtubule network despite containing the microtubule-binding region^15^. In theory, this identifies an alternative therapeutic approach for patients with the longer variants through use of HSP90 inhibitors, although to date clinical trials of HSP90 inhibitors have not shown efficacy in ALK-positive patients with these variants^18, 19, 20^. Moreover, HSP90 inhibitors would not be useful in patients with EML4-ALK V3a, V3b or V5 as these fusion proteins are not dependent upon HSP90 for their expression. Furthermore, patients with V3a and V3b respond less well to ALK inhibitor treatment suggesting additional tumorigenic mechanisms that may be independent of ALK kinase activity^12^. This is important as, although V5 is rare, V3a and V3b represent up to 50% of EML4-ALK fusions in NSCLC^8, 12, 14, 21, 22^.

Wild-type EML proteins have been found to interact with members of the human NEK kinase family^23^. The human genome encodes eleven NEKs, named Nek1 to Nek11, many of which exhibit cell cycle-dependent activity and have functions in microtubule organization^24, 25, 26^. NEK9, NEK6 and NEK7 are activated in mitosis and act in concert to regulate mitotic spindle assembly with NEK9 upstream of NEK6 and NEK7^27^. NEK6 and NEK7 are the shortest of the NEKs consisting solely of a kinase domain with a short N-terminal extension. Moreover, they are closely related being ∼85% identical within their catalytic domains. In contrast, NEK9 is one of the longest NEKs, consisting of an N-terminal catalytic domain (residues 52-308) followed by a C-terminal regulatory region (residues 309-979). Within this C-terminal region is an RCC1 (regulator of chromatin condensation 1)-like domain (residues 347-726) predicted to fold into a seven-bladed β-propeller. Biochemical studies indicate that deletion of the RCC1-like domain causes constitutive activation of NEK9, indicating that this acts as an auto-inhibitory domain^28^. This domain is followed by a C-terminal tail that contains a coiled-coil motif (residues 891-939) that promotes NEK9 dimerization^28^. NEK9 directly interacts with NEK7, and presumably NEK6, through a sequence that lies between the RCC1-like domain and coiled-coil^29^. NEK9 stimulates NEK6 and NEK7 kinase activity by multiple mechanisms, including activation loop phosphorylation, dimerization-induced autophosphorylation, and allosteric binding-induced reorganization of catalytic site residues^29, 30, 31^.

Here, we demonstrate that expression of not only activated NEK9 and NEK7 kinases but also full-length EML4 and the short EML4-ALK variants that bind microtubules (V3 and V5) alters cell morphology, disturbs cell polarity and promotes cell migration. These changes are not seen in cells expressing the longer variants (V1 and V2) that do not bind microtubules. Moreover, our data reveal interaction between the EML4 NTD and NEK9, and suggest a model in which the EML4-ALK V3 and V5 proteins stabilise microtubules and perturb cell behaviour through recruitment of NEK9 and NEK7. Finally, we show that EML4-ALK lung cancer patients exhibit a significant correlation between expression of V3/V5 and high levels of NEK9 protein, and that elevated NEK9 expression is also associated with worse progression-free and overall survival raising the prospect of novel therapeutic approaches.

## RESULTS

### Constitutively active NEK9 leads to microtubule-dependent changes in interphase cell morphology

We initially set out to explore the functional consequences of untimely activation of NEK9 in interphase cells. For this, an inducible system was established in which expression of myc-tagged wild-type NEK9, a full-length kinase-inactive mutant (K81M), and a constitutively-activated mutant lacking the RCC1-like domain (ΔRCC1) was placed under the control of a tetracycline-inducible promoter in U2OS osteosarcoma cells (Fig. 1A). Western blots indicated time-dependent induction of expression of all three recombinant proteins upon addition of doxycycline, while subcellular localization confirmed the previously described cytoplasmic localization of the full-length and activated (ΔRCC1) proteins, and nuclear localization of the catalytically-inactive mutant^28^ (Fig. 1B; and Supp. Fig. S1A). Unexpectedly, brightfield and immunofluorescence microscopy revealed that induction of activated NEK9 led to a change in interphase cell morphology with formation of long cytoplasmic protrusions that were not seen upon induction of either wild-type or catalytically-inactive NEK9 (Fig. 1C-E; and Supp. Fig. S1B). Flow cytometry revealed no significant change in cell cycle distribution upon induction of these proteins (Supp. Fig. S1C). Depolymerisation of the microtubule network with nocodazole or vinorelbine led to loss of these protrusions, while time-lapse imaging indicated that these protrusions were comparatively stable structures that did not undergo retraction except during cell division (Fig. 1F, G; and Supp. Mov. S1, S2). Super-resolution radial fluctuations (SRRF) microscopy^32^ revealed that these protrusions contained not only microtubules and the activated NEK9 protein, but also actin and acetylated tubulin (Fig. 1H, I). Indeed, Western blotting indicated that induction of activated NEK9 was associated with an increased abundance in cells of acetylated tubulin, a marker of stabilised microtubules (Fig. 1J). Consistent with this, time-lapse imaging of live cells incubated with the fluorescent SiR-tubulin probe^33^ demonstrated that microtubules were more resistant to nocodazole-induced depolymerization following induction of the activated NEK9 kinase (Fig. 1K; and Supp. Fig. S1D). Although the mechanism for microtubule stabilization remains to be determined, we found that inhibition of the microtubule-based kinesin motor Eg5 with STLC also blocked this morphological change (Fig. 1F), in line with recent reports that Eg5 promotes microtubule stabilization in post-mitotic neurons^34^. Hence, activation of NEK9 leads to a microtubule-dependent change in interphase cell morphology characterised by formation of elongated cytoplasmic protrusions that contain both actin and stabilised microtubules.

**Figure 1.**
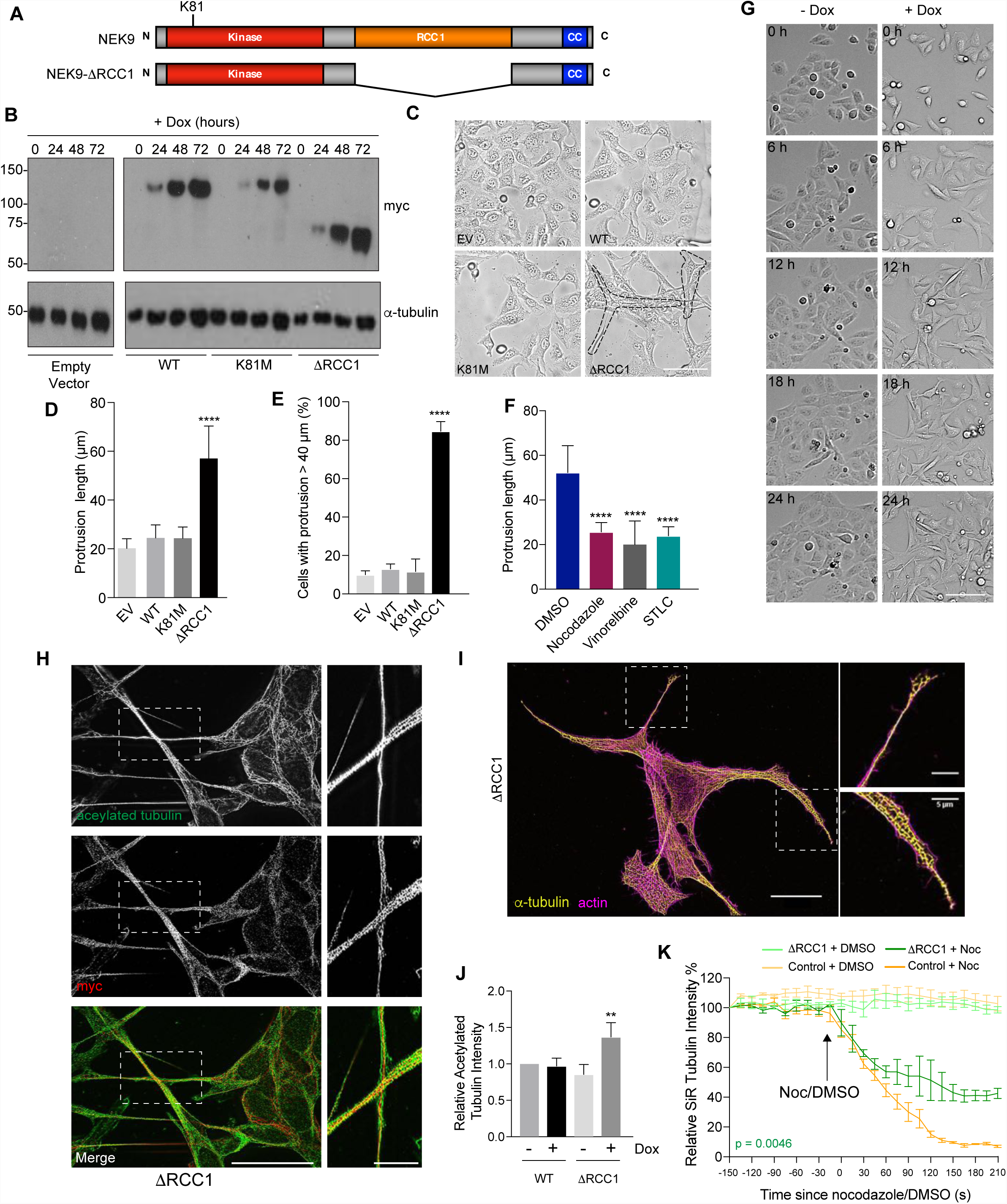
Constitutively active NEK9 induces altered morphology and stabilised microtubules in interphase cells. **A**. Schematic representation of full-length and activated NEK9 indicating the kinase, RCC1 and coiled-coil (CC) domains. The position of the inactivating K81M mutation is indicated. **B**. Lysates were prepared from U2OS stable cell lines induced to express wild-type (WT), catalytically-inactive (K81M) or constitutively-active (ΔRCC1) myc-NEK9 with doxycycline for 0, 24, 48 and 72 h, and analysed by Western blot with myc and α-tubulin antibodies. U2OS cells with the tetracycline inducible plasmid (empty vector, EV) were used as a negative control. M. wts (kDa) are indicated on the left. **C**. U2OS cells were induced to express constructs as in B for 48 h prior to imaging by phase contrast microscopy; scale bar, 50 µm. **D**. The maximum length of cytoplasmic protrusions, measured from nucleus to the most distant cytoplasmic point, for cells shown in C is indicated. **E**. The percentage of cells with protrusions exceeding 40 µm for cells in C & D is indicated. **F**. U2OS cells induced to express myc-NEK9-ΔRCC1 for 72 h were treated for 4 h with DMSO, nocodazole, vinorelbine or STLC, and then analysed by phase-contrast microscopy; the maximum length of cytoplasmic protrusions measured. **G**. Stills from time-lapse phase-contrast imaging of U2OS:myc-NEK9-ΔRCC1 cells treated with or without doxycycline for the times indicated; scale bar, 100 µm. **H**. U2OS:myc-NEK9-ΔRCC1 cells treated with doxycycline for 48 h were analysed by immunofluorescence microscopy with myc and α-tubulin antibodies and images reconstructed using SRRF analysis. **I**. U2OS:myc-NEK9-ΔRCC1 cells treated with doxycycline for 48 h were analysed by immunofluorescence microscopy with α-tubulin antibodies while actin was stained with phalloidin-FITC. Images were reconstructed using SRRF analysis. In H & I, magnified views of cytoplasmic protrusions are shown on the right from the boxed regions in cells on the left. Scale bars in the main images, 20 µm, and in the zooms, 5 µm. **J**. Cell lysates prepared from U2OS:myc-NEK9-WT and ΔRCC1 cells +/- 48 h doxycycline treatment were Western blotted with acetylated tubulin and GAPDH antibodies. The intensity of acetylated tubulin relative to GAPDH is indicated. **K**. U2OS:myc-NEK9-WT and ΔRCC1 cells +/- 48 h doxycycline treatment were incubated with SiR-Tubulin to visualise microtubules and SiR-Tubulin intensity measured every 15 s following addition of nocodazole or DMSO. Data represent means from 3 separate experiments with 8 wells per condition.

### Activated NEK9 perturbs interphase cell polarity and stimulates cell migration

The morphological changes observed upon expression of activated NEK9 was characteristic of that seen in migrating cells. To test whether activated NEK9 also altered cell migration, the rate of closure of a scratch wound was measured. This revealed that cells induced to express the activated NEK9 closed the wound more quickly than cells containing an empty vector treated with doxycycline or cells containing the activated NEK9 construct but without induction (Fig. 2A, B). Surprisingly, this was not due to increased persistence in directional migration as these cells migrated with reduced straightness compared to control cells (Fig. 2C). Indeed, cells at the wound edge failed to orient either their Golgi network or their centrosomes towards the wound edge in the presence of activated NEK9 suggestive of loss of cell polarity (Fig. 2D-F). Isolated cells also exhibited disruption of the Golgi network that was dispersed over a larger area in cells expressing activated NEK9 (Fig. 2G, H). Individual cell tracking experiments confirmed a significant increase in the velocity of migration as well the distance travelled in isolated cells induced to express activated NEK9 (Fig. 2I-K). Furthermore, there was a decrease in the straightness of migration in isolated cells in the presence of activated NEK9 consistent with more frequent changes in direction of migration (Fig. 2L). The wide variance in the comparator groups likely reflects the stochastic nature of migration of isolated cells in culture that may be influenced by factors such as cell cycle stage and cell density. Finally, continuous monitoring of cell migration using a real-time transwell migration assay (xCELLigence Real Time Cell Assay) not only provided additional evidence that induction of activated NEK9 led to a significant increase in cell migration but also demonstrated that this was evident at 10 hours after induction and is therefore is not a result of increased proliferation (Fig. 2M, N). Taken together, these data demonstrate that expression of activated NEK9 not only leads to a change in interphase cell morphology but also perturbs cell polarity and enhances cell migration.

**Figure 2.**
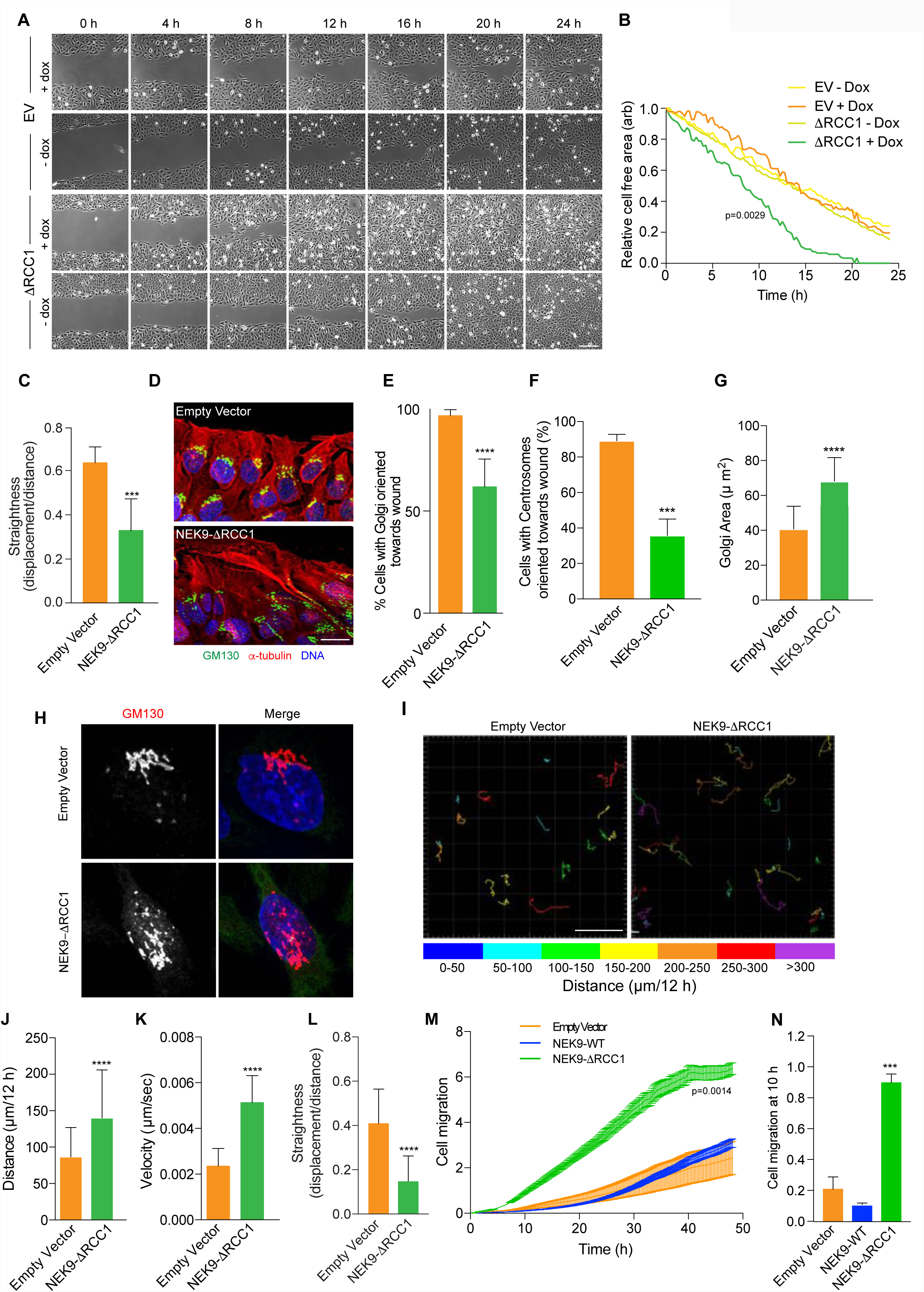
Activated NEK9 disturbs cell polarity and increases cell migration. **A**. U2OS:myc-NEK9-ΔRCC1 or -EV (empty vector) cells were induced with or without doxycycline for 48 h before time-lapse imaging was used to observe closure of a scratch wound. Representative images at the time-points indicated are shown; scale bar, 200 µm. **B**. The rate of wound closure for cells in A is shown. Data represent means from 3 separate experiments with 8 positions measured in each treatment. **C**. The persistence of movement for cells at the leading edged of the wound generated as in A was measured by quantifying the straightness of track for each cell. **D**. Cells were treated as in A for 6 h before being analysed by immunofluorescence microscopy with antibodies against the Golgi marker, GM130 (green), and α-tubulin (red). DNA (blue) was stained with Hoechst 33258; scale bar, 20 µm. **E**. The orientation of the Golgi network with respect to the wound was scored for each cell in the front row of cells migrating into the wound. **F**. Cells treated as in A were analysed by immunofluorescence microscopy with GM130 and γ-tubulin antibodies. Orientation of centrosomes with respect to the wound was scored for each cell in the front row of cells migrating into the wound. **G**. U2OS:myc-NEK9-ΔRCC1 or EV were induced for 48 h prior to being analysed by immunofluorescence microscopy with GM130 antibodies. The area occupied by the Golgi network was measured. **H**. Cells treated and stained as in G; DNA was stained with Hoechst 33258. Scale bar, 10 µm. **I**. U2OS:myc-NEK9-ΔRCC1 or –EV were induced for 24 h prior to being transfected with YFP protein for 24 h and subsequent time-lapse imaging. The movement of individual cells was tracked over a 12 h period by following the position of the YFP signal; distance travelled is colour-coded as indicated (µm). Scale bar, 200 µm. **J**. The mean distance travelled by cells treated as in E is indicated. **K**. The mean velocity of cells treated as in E is indicated. **L**. The persistence of movement for cells in E was measured by quantifying the straightness of track (track displacement/track length) for each cell. **M**. U2OS:myc-NEK9-WT, -ΔRCC1 or -EV were induced for 24 h before being starved in serum-free medium for a further 24 h, and then seeded in RTCA CIM-16 plates. Complete medium containing 10% FBS was used as a chemo-attractant and impedance measurements were used to determine cell migration in real time. Data represent means and S.D. from 3 separate experiments. **N**. The histogram shows the cell migration index at 10 h from the RTCA assays presented in M.

### NEK9-induced changes in cell morphology and migration are dependent on NEK7

A major function of NEK9 activity in mitosis is to activate NEK6 and NEK7^24^. This is thought to occur via both phosphorylation of activation loop residues and allosteric binding^30, 31^. Indeed, biochemical and structural studies have revealed direct interaction between residues 810-828 of NEK9 and a groove on the C-terminal lobe of the NEK7 catalytic domain^29^. To determine whether the change in interphase cell morphology and migration induced by activated NEK9 is dependent on NEK6 or NEK7, cells expressing activated NEK9 were depleted of NEK6 or NEK7 (Supp. Fig. S2A). Quantitative imaging revealed that, in contrast to mock-or NEK6-depleted cells that still exhibited extended cytoplasmic protrusions upon induction of activated NEK9, depletion of NEK7 prevented formation of these protrusions (Fig. 3A; and Supp. Fig. S2B). This suggested that this phenotype was dependent upon the untimely activation of NEK7 by NEK9. To confirm this hypothesis, we made use of a constitutively active mutant version of NEK7. Previous studies had revealed that, in the absence of NEK9, NEK7 adopts an auto-inhibited conformation in which the side-chain of Tyrosine-97 blocks the active site and that mutation of this residue to alanine relieves this inhibition^31^. Strikingly, transient expression of the activated NEK7-Y97A mutant, but not wild-type NEK7, caused generation of similar elongated cytoplasmic protrusions in U2OS cells (Fig. 3B, C). Consistent with the lack of effect of NEK6 depletion, expression of the equivalent activated NEK6-Y108A mutant did not cause formation of cytoplasmic protrusions but rather caused cells to become more rounded. Real-time transwell assays indicated that the increase in cell migration induced by activated NEK9 was also dependent on NEK7 but not NEK6 and, as it was detected after 10 hours, unlikely to be explained by reduced proliferation (Fig. 3D, E; and Supp. Figs. S2C, D).

**Figure 3.**
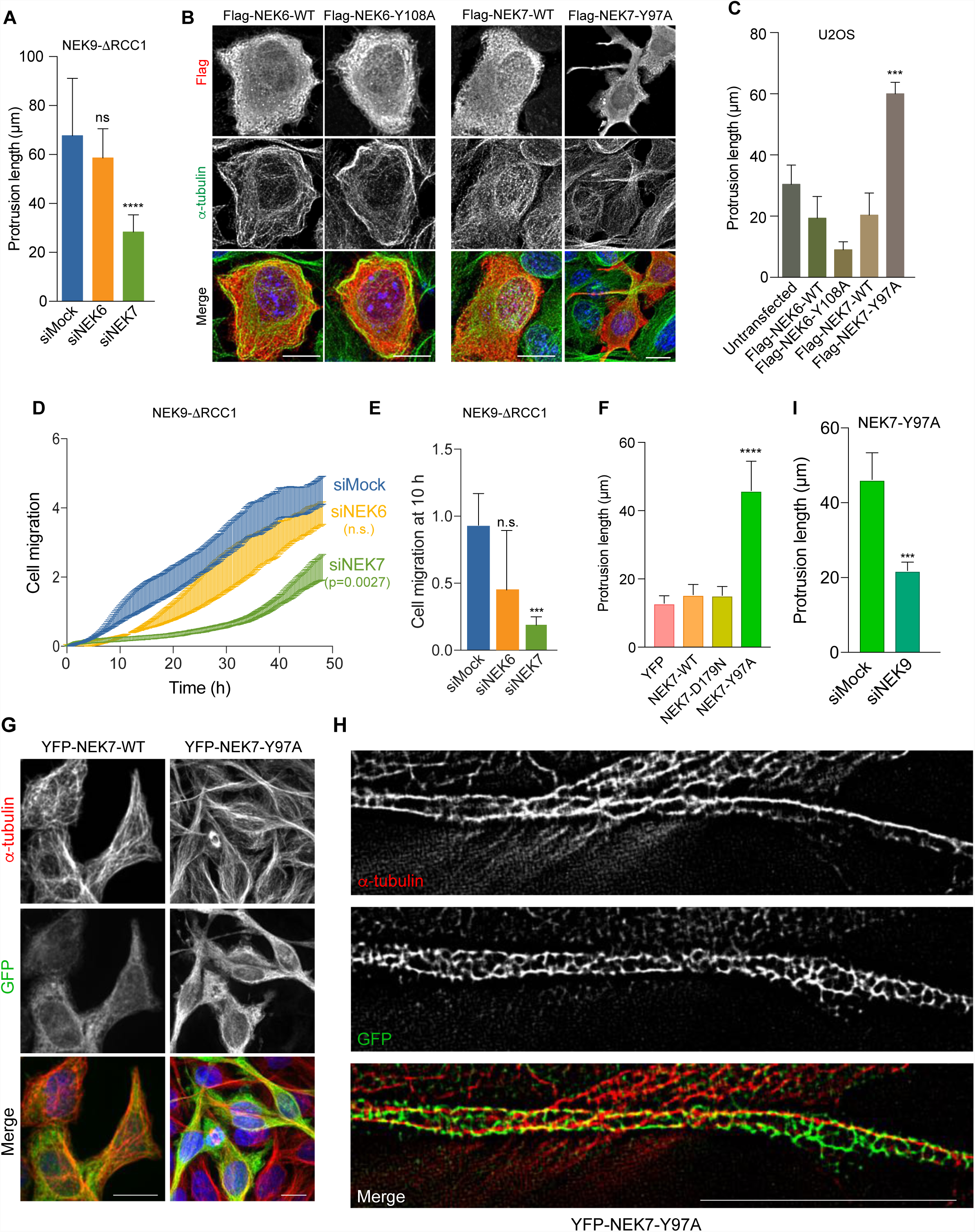
NEK9-induced alterations in cell morphology and migration are dependent on NEK7. **A**. U2OS:myc-NEK9-ΔRCC1 cells were depleted with mock, NEK6 or NEK7 siRNAs for 24 h prior to being induced with doxycycline for a further 48 h. Cells were then processed for immunofluorescence microscopy and the maximum length of interphase cytoplasmic protrusions measured. **B**. U2OS cells were transiently transfected with wild type (WT) or constitutively active Flag-NEK6 (Y108A) or Flag-NEK7 (Y97A), as indicated, for 24 h before being processed for immunofluorescence microscopy with Flag (green) and α-tubulin (red) antibodies. DNA (blue in merge) was stained with Hoechst 33258; scale bars, 10 µm. **C**. The maximum length of interphase cytoplasmic protrusions for cells in B was measured. **D**. U2OS:myc-NEK9-ΔRCC1 cells were depleted with mock, NEK6 or NEK7 siRNAs for 24 h prior to being induced with doxycycline for a further 48 h and cell migration analysed in real time as in Fig. 2I. Data represent means and S.D. from 3 separate experiments. **E**. The histogram shows the cell migration index at 10 h from the RTCA assays presented in D. **F**. HeLa:YFP alone, or YFP-NEK7-WT, D179N or Y97A were induced with doxycycline for 72 h before being processed for immunofluorescence microscopy and the maximum length of interphase cytoplasmic protrusions measured. **G**. HeLa:YFP-NEK7-WT and Y97A cells treated with doxycycline for 48 h were analysed by immunofluorescence microscopy with GFP (green) and α-tubulin (red) antibodies. Scale bar, 20 µm. **H**. HeLa-YFP-NEK7-Y97A cells were treated as in H and images generated using super-resolution reconstruction microscopy. Scale bar, 5 µm. **I.** HeLa:YFP-NEK7-Y97A were depleted with mock or NEK9 siRNAs for 24 h prior to being induced with doxycycline for a further 48 h. Cells were then processed for immunofluorescence microscopy and the maximum length of interphase cytoplasmic protrusions measured.

To further address the role of NEK7 in this pathway, we established HeLa cell lines in which YFP tagged wild-type, catalytically-inactive (D179N) and constitutively-active (Y97A) versions of NEK7 were expressed under the control of a tetracycline inducible promoter (Fig. S3). Consistent with transient expression, inducible expression of activated NEK7 but not wild-type or kinase-inactive NEK7 resulted in the formation of microtubule rich cytoplasmic protrusions in interphase (Fig. 3F, G). Analysis of the localization of the activated NEK7 protein by SRRF microscopy revealed substantial co-localisation with microtubules in the cytoplasmic protrusions (Fig. 3H). Interestingly, these protrusions formed upon expression of activated NEK7 were lost upon depletion of NEK9 suggesting a more complex relationship than NEK9 simply acting upstream to enhance NEK7 catalytic activity (Fig. 3I). Together, these data indicate that the microtubule-dependent alteration of cell morphology and increased rate of migration that occur upon activation of NEK9 is dependent on NEK7, but equally that similar changes induced upon activation of NEK7 are dependent on NEK9.

### EML4 interacts with NEK9 and promotes cell migration

To determine the mechanism through which the NEK9-NEK7 pathway may cause this change in cell morphology and migration, myc-tagged wild-type NEK9 was immunoprecipitated from the stable cell line and immune complexes analysed by mass spectrometry. Intriguingly, this identified not only tubulin but also the microtubule-associated protein, EML4, as a NEK9 binding partner (Supp. Fig. S4A-C). Previous large-scale proteomic mapping had identified EML2, EML3 and EML4 as potential binding partners of NEK6^23^, while we have found that both EML3 and EML4 can be phosphorylated in vitro by NEK6 and NEK7 (R.A., L.O., M.W.R., R.B., A.M.F., manuscript in preparation). Western blotting of immunoprecipitates prepared with antibodies against either NEK9 or EML4 confirmed interaction between endogenous EML4 and NEK9 in extracts prepared from asynchronous U2OS cells (Fig. 4A). Moreover, YFP-EML4 was found to co-precipitate equally well with the activated NEK9-ΔRCC1 mutant as wild-type NEK9 (Fig. 4B). NEK9 is also known to bind directly through its C-terminal region to NEK7 and NEK6, an interaction that is normally blocked in interphase by competitive interaction with the LC-8 adaptor protein^29, 35^. Removal of the RCC1 domain relieves the auto-inhibited conformation and leads to activation of NEK9^28^. This in turn enables autophosphorylation at Ser-944, which displaces LC-8 and allows interaction with NEK7 and NEK6^35^. Hence, we propose that activated NEK9 can potentially interact simultaneously with both EML4 and NEK7.

**Figure 4.**
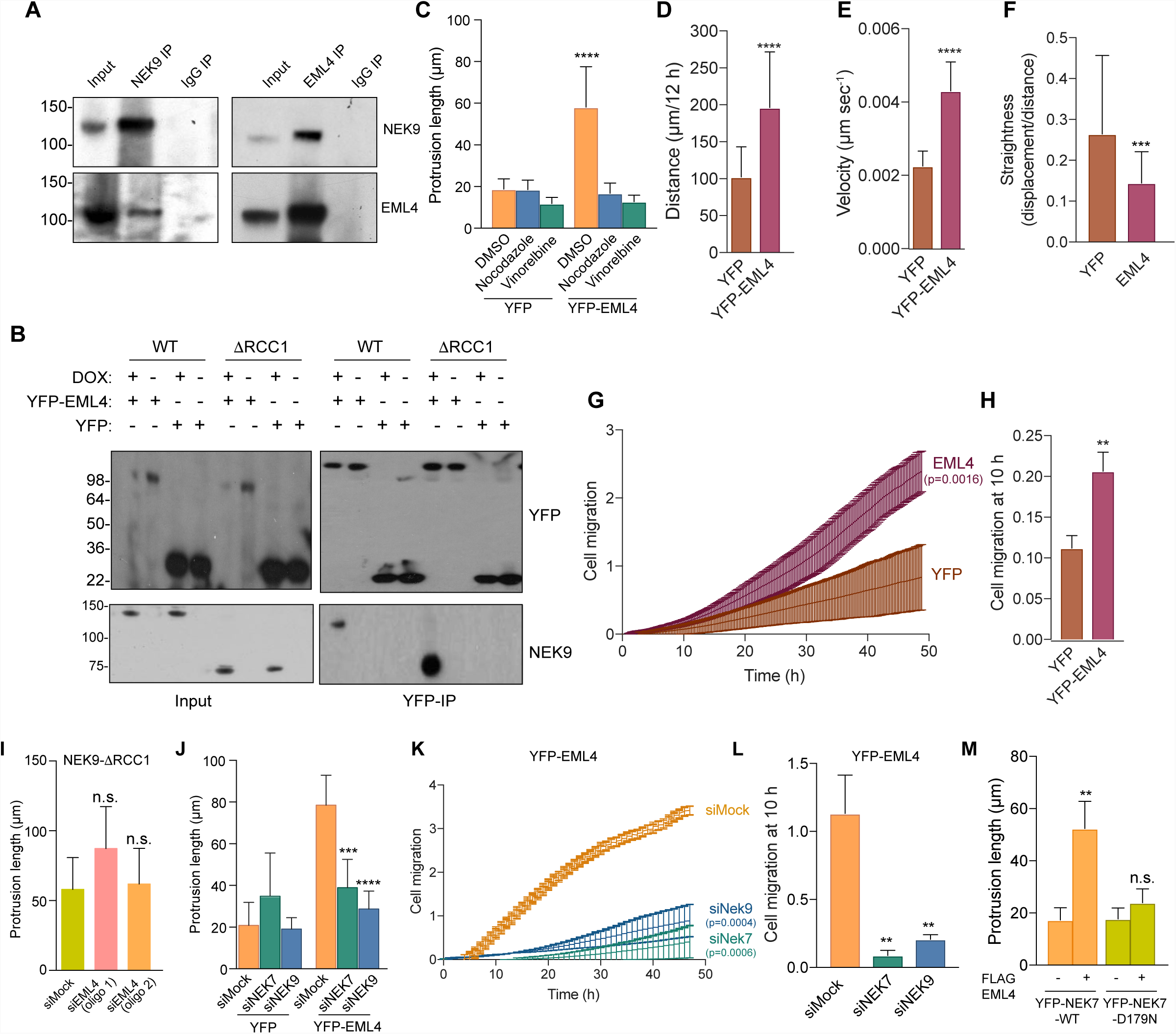
EML4 induces NEK9-dependent changes in cell morphology and migration. **A**. U2OS cell lysates (inputs) and immunoprecipitates (IP) prepared with rabbit NEK9 and control IgGs were analysed by Western blot with the antibodies indicated. M. wts (kDa) are indicated on the left. **B**. U2OS:myc-NEK9-WT or ΔRCC1 cells were induced for 48 h before being transfected with YFP-EML4 or YFP alone for a further 24 h. Lysates (input) and Immunoprecipitates (IP) were prepared with GFP antibodies and analysed by Western blot with the antibodies indicated. **C**. U2OS:YFP or U2OS:YFP-EML4 cell lines were treated with DMSO, nocodazole or vinorelbine for 16 h as indicated before analysis by phase contrast microscopy and the maximum length of interphase cytoplasmic protrusions measured. **D**. Individual cell tracking experiments were undertaken with U2OS:YFP and U2OS:YFP-EML4 cell lines and analysed as in Fig. 2E. The mean distance travelled is indicated. **E**. The mean velocity of cells treated as in D is indicated. **F**. The track straightness of cells treated as in D is indicated. **G**. U2OS:YFP and U2OS:YFP-EML4 cell lines were starved in serum-free medium for 24 h, and cell migration analysed in real time as in Fig. 2M. Data represent means and S.D. from 3 separate experiments. **H**. The histogram shows the cell migration index at 10 h from the data in G. **I**. U2OS:myc-NEK9-ΔRCC1 were depleted with mock or two different siRNAs against EML4 and the maximum length of interphase cytoplasmic protrusions measured. **J**. U2OS:YFP or U2OS:YFP-EML4 cells were mock-, NEK7-or NEK9- depleted for 24 h before being analysed by phase contrast microscopy and the maximum length of interphase cytoplasmic protrusions measured. **K**. U2OS:YFP-EML4 cells were mock-, NEK9-or NEK7-depleted for 48 h, starved in serum-free medium for a further 24 h, and cell migration analysed in real time as in Fig. 2M. Data represent means and S.D. from 3 separate experiments. **L**. The histogram shows the cell migration index at 10 h from the data in K. **M**. HeLa:YFP-NEK7-WT or D179N were induced for 24 h with doxycycline before being transfected with Flag-EML4 for a further 24 h and maximum length of cytoplasmic protrusions for transfected cells measured.

To investigate whether the microtubule-dependent changes in cell morphology that result from expression of activated NEK9 and NEK7 may relate to its interaction with EML4, U2OS stable cell lines were generated that expressed YFP alone or YFP-EML4. These cell lines expressed tagged proteins of the predicted size as determined by Western blotting while the YFP-EML4 protein localized as expected to the microtubule network (Supp. Fig. S5A-C). In contrast to parental cells or cells expressing YFP alone, the YFP-EML4 cells exhibited a morphology similar to cells expressing activated NEK9 or NEK7 with extended cytoplasmic protrusions that were lost upon treatment with the microtubule depolymerizing agents, nocodazole or vinorelbine (Fig. 4C; and Supp. Fig. S5D). Individual cell tracking experiments revealed expression of YFP-EML4 also led to as increased distance and velocity of migration, but reduced straightness, compared to those expressing YFP alone (Fig. 4D-F). Real-time transwell migration assays confirmed that YFP-EML4 expression led to an increased rate of migration as compared to YFP alone (Fig. 4G, H). To test the relationship of EML4 and NEK9 in the generation of these phenotypes, these proteins were depleted in the different stable cell lines. Depletion of EML4 did not prevent the alterations in cell morphology induced upon expression of activated NEK9 (Fig. 4I; and Supp. Fig. S5E). In contrast, depletion of NEK9 or NEK7 led to loss of cytoplasmic protrusions and reduced migration in cells expressing YFP-EML4 (Fig. 4J-L; and Supp. Fig. S5F, G). Furthermore, induced expression of catalytically-inactive NEK7 prevented formation of cytoplasmic protrusions in cells overexpressing EML4 (Fig. 4M). These data indicate that overexpression of EML4 induces altered cell morphology and enhanced migration that is dependent upon NEK9 and NEK7, and can be blocked by expression of inactive NEK7.

### EML4 recruits NEK9 and NEK7 to microtubules

To understand how NEK9 and NEK7 act downstream of EML4 to induce alterations in cell morphology and migration, we first examined which region of the EML4 protein may be responsible for interaction with NEK9. Co-immunoprecipitation experiments revealed that endogenous NEK9 could interact not only with full-length EML4 but also both the isolated NTD and TAPE domains (Fig. 5A). On the other hand, it did not interact with a shorter fragment of the NTD that only encompassed the trimerization domain. As NEK9 could interact with the microtubule-binding EML4 NTD, we hypothesized that EML4 may promote recruitment of NEK9 and NEK7 to the microtubule cytoskeleton. To test this hypothesis, we examined the localization of endogenous NEK9 in cells expressing different EML4 constructs. In cells expressing the TAPE domain, endogenous NEK9 was mainly distributed in the cytoplasm with only faint detection on microtubules, whereas in cells expressing the NTD, NEK9 exhibited strong recruitment to microtubules (Fig. 5B, C; and Supp. Fig. S6A). Further studies using the stable HeLa:YFP-NEK7 cell line revealed that YFP-NEK7 was recruited to microtubules upon transfection of Flag-tagged EML4 NTD, but not the TAPE domain (Fig. 5D, E; and Supp. Fig. S6B). Together, these data suggest a model in which the EML4 NTD recruits both NEK9 and NEK7 to microtubules. This explains not only how EML4 can act upstream of NEK9 and NEK7, but also why the NEK9 protein is required for activated NEK7 to promote microtubule-dependent changes in cell morphology and migration.

**Figure 5.**
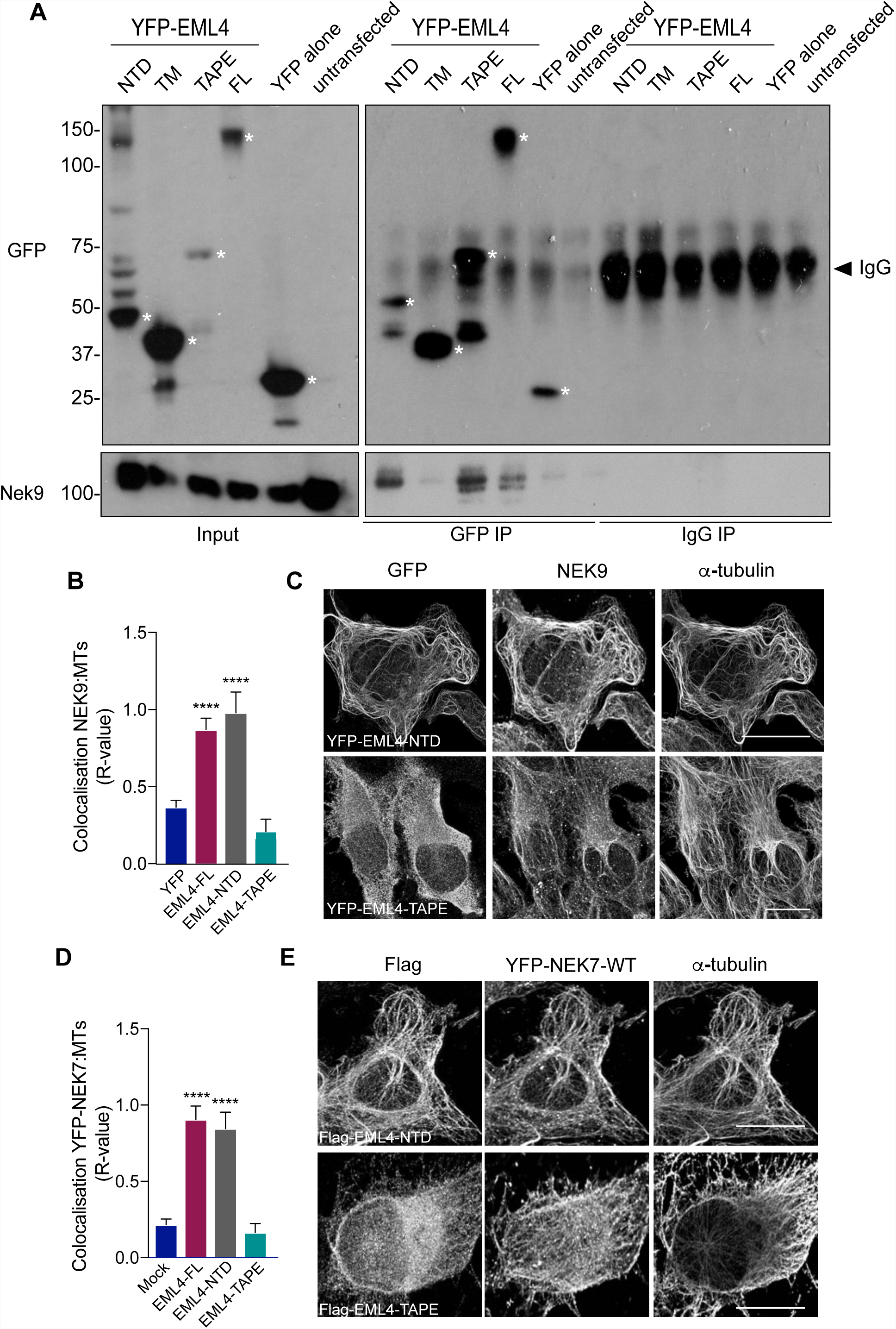
EML4 recruits NEK9 and NEK7 to interphase microtubules. **A**. U2OS cells were transfected with YFP-EML4-full length (FL), TM (trimerisation motif), TAPE, NTD or YFP alone for 24 h. Lysates (inputs) and immunoprecipitates (IP) were prepared with GFP and control IgGs and analysed by Western blot with the antibodies indicated. Asterisks indicate the proteins of interest. **B**. U2OS cells were transfected with YFP-EML4 N-terminal domain (NTD) or TAPE domain (TAPE), as indicated, for 24 h before being processed for immunofluorescence microscopy with antibodies against GFP, NEK9 and α-tubulin, and colocalization between the Nek9 and α-tubulin signal calculated. **C.** Representative images of cells in B. Scale bar, 10 µm. **D**. HeLa:YFP-NEK7 cells were induced for 48 h with doxycycline before being mock transfected or transfected with Flag-EML4-NTD or TAPE for a further 24 h. Cells were then processed for immunofluorescence microscopy with antibodies against Flag, GFP and α-tubulin. Co-localization between the YFP-NEK7 fluorescence and α-tubulin signal was calculated. **E**. Representative images of cells in D. *R* values in B and D show the mean Pearson’s correlation coefficient from 5 lines per cell in 10 cells (±S.D.).

### EML4-ALK variants that localise to microtubules induce altered cell morphology, polarity and migration

Considering their importance in cancer biology, we were intrigued to know whether EML4-ALK fusion proteins might also alter cell morphology and migration. To explore this possibility, we generated U2OS cell lines with stable expression of the ALK kinase domain alone or four of the different EML4-ALK variants (Fig. 6A). These included the longer variants, V1 and V2, which encode unstable proteins that do not localise to microtubules, and the shorter variants, V3a and V5, which encode stable proteins that localise to microtubules^15, 16^. Strikingly, microscopic analysis revealed that those cells in which the variant localised to interphase microtubules (V3a and V5) formed extended cytoplasmic protrusions, whereas those cells in which the variant did not localise to the microtubules (V1 and V2) did not (Fig. 6B-D). Furthermore, both individual cell tracking and real-time transwell assays revealed that expression of variants 3a and 5 stimulated cell migration with significant increases in distance and velocity and decreased straightness, whereas expression of V1 and V2 did not (Fig. 6E-I). To assess the consequences of these variants on cell polarity, Golgi orientation was measured in a wound-healing assay with U2OS cells expressing either EML4-ALK V1 or V3a. The percentage of cells in which the Golgi was oriented towards the wound was lower in cells expressing V1 (75%) than in parental U2OS cells, potentially due to a modest effect on microtubule organization. However, there was a significant reduction in Golgi orientation in cells expressing V3a (58%) as compared to V1 (Fig. 6J). We also analysed centrosome orientation and found that cells expressing V3a had substantially reduced ability to orient their centrosomes towards the wound edge, while single cell tracking of cells at the wound edge revealed a significant decrease in straightness of migration (Fig. 6K & L). Furthermore, the area occupied by the Golgi network was significantly greater in isolated cells expressing V3a as compared to ALK alone or V1 (Fig. 6M & N). Hence, expression of the short EML4-ALK variants that localize to microtubules lead to altered morphology, reduced polarity and enhanced migration as compared to cells expressing the longer EML4-ALK variants that do not localise to microtubules.

**Figure 6.**
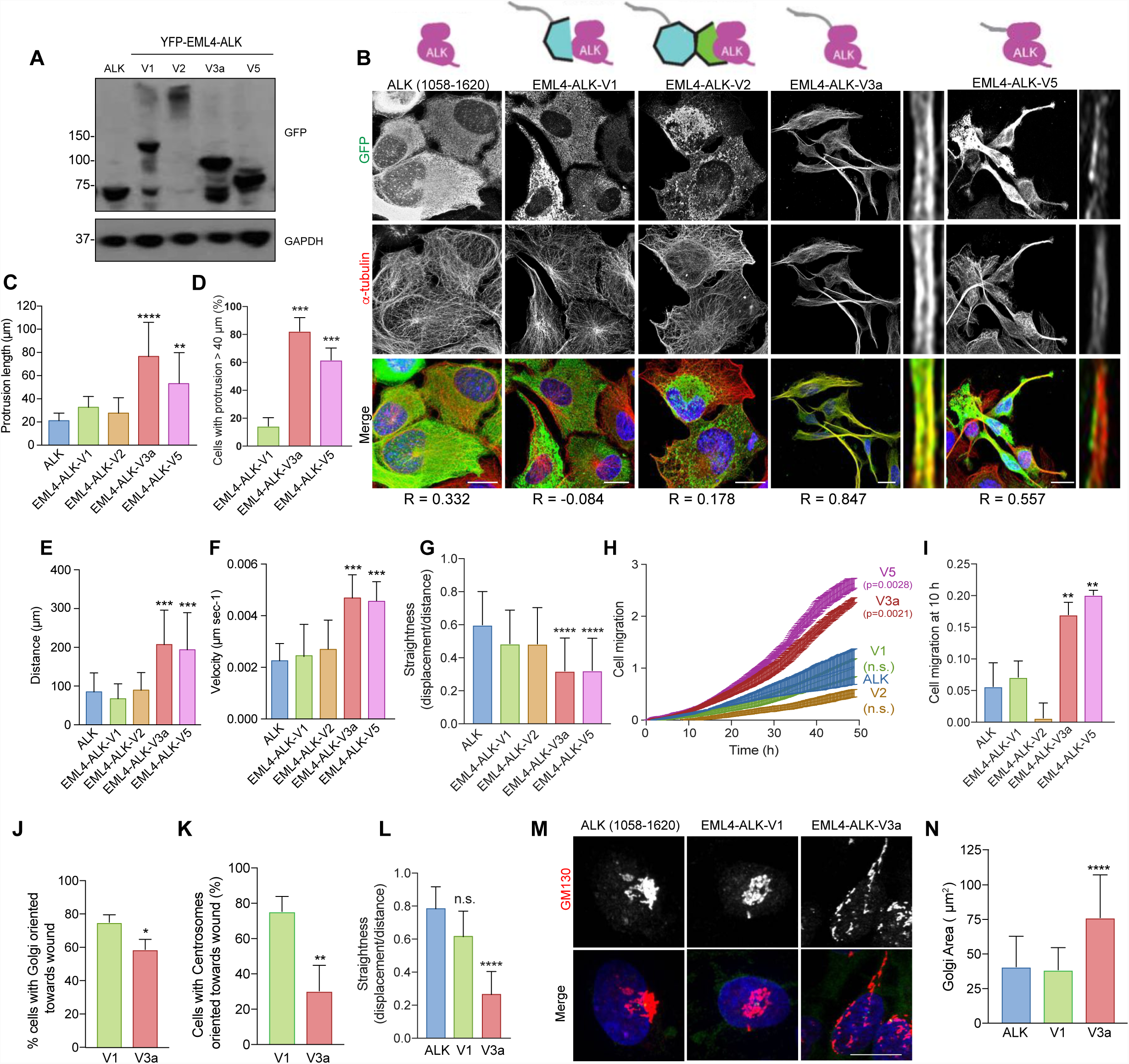
EML4-ALK variants that bind microtubules alter cell morphology and migration. **A.** U2OS cell lines stably expressing YFP-tagged EML4-ALK fusion variants or the ALK kinase domain alone were lysed and analysed by Western blot with the antibodies indicated. M. wts (kDa) are indicated on the left. **B**. U2OS stable cell lines expressing YFP-tagged proteins as in A were analysed by immunofluorescence microscopy with antibodies against GFP (green) and α-tubulin (red). DNA (blue) was stained with Hoechst 33258 (blue in merge). Magnified views of individual microtubules are shown on the right of V3a and V5 images shown on the left. Schematic versions of the YFP-tagged proteins and the Pearson’s correlation coefficient (*R*) of co-localization with microtubules are shown. Scale bars, 10 µm. **C**. The maximum length of interphase cytoplasmic protrusions for cells in B was measured. **D**. The % of cells from B and C with protrusions greater than 40 µm is indicated. **E**. Individual cell tracking experiments were undertaken with cell lines as in A and analysed as in Fig. 2I. The mean distance travelled is indicated. **F.** As for E, but showing mean velocity of movement. **G**. As for E but showing track straightness. **H**. Cell lines as described in A were incubated in serum-free medium for 24 h and cell migration analysed in real time as in Fig. 2M. **I**. Histogram shows the cell migration index at 10 h from the data in H. **J**. U2OS:YFP-EML4-ALK-V1 or V3a cells were grown until confluence before a scratch wound was made in the monolayer. Cells were analysed after 6 h by immunofluorescence microscopy with antibodies against the Golgi marker, GM130 (green), and the orientation of the Golgi network with respect to the wound scored for each cell at the wound edge. **K**. Cells were treated as in J and analysed by immunofluorescence microscopy with γ-tubulin antibodies. The orientation of centrosomes with respect to the wound was scored for each cell at the wound edge. **L**. Cells were treated as in J prior to time-lapse imaging of wound closure. The persistence of movement for cells was measured by quantifying the straightness of track for each cell at the leading edged of the wound. **M**. Isolated U2OS:YFP-EML4-ALK-V1 or V3a cells were analysed by immunofluorescence microscopy for GM130 antibodies (red). DNA was stained with Hoechst 33258. Scale bar, 10 µm. **N**. The area occupied by the Golgi network in cells in M was measured.

### Changes mediated by EML4-ALK V3 depend on NEK9 and NEK7 but not ALK activity

To determine whether the changes observed in cell morphology and migration induced by EML4-ALK V3 required catalytic activity of the ALK tyrosine kinase, we first generated U2OS stable cell lines with catalytically-inactive (D1270N) versions of EML4-ALK V1 and V3. This revealed that cells expressing the inactive mutant proteins had similar length cytoplasmic protrusions as those seen in cells expressing the wild-type proteins (Fig. 7A). Second, we determined the consequences on cell migration of treatment with the ALK inhibitor, crizotinib. This did not alter the rate of migration of U2OS cells expressing EML4-ALK V3a despite preventing ALK-dependent signalling in these cells as expected (Fig. 7B, C; and Supp. Fig. S7A). Hence, we conclude that the altered morphology and enhanced migration of cells expressing EML4-ALK V3 does not require ALK activity.

**Figure 7.**
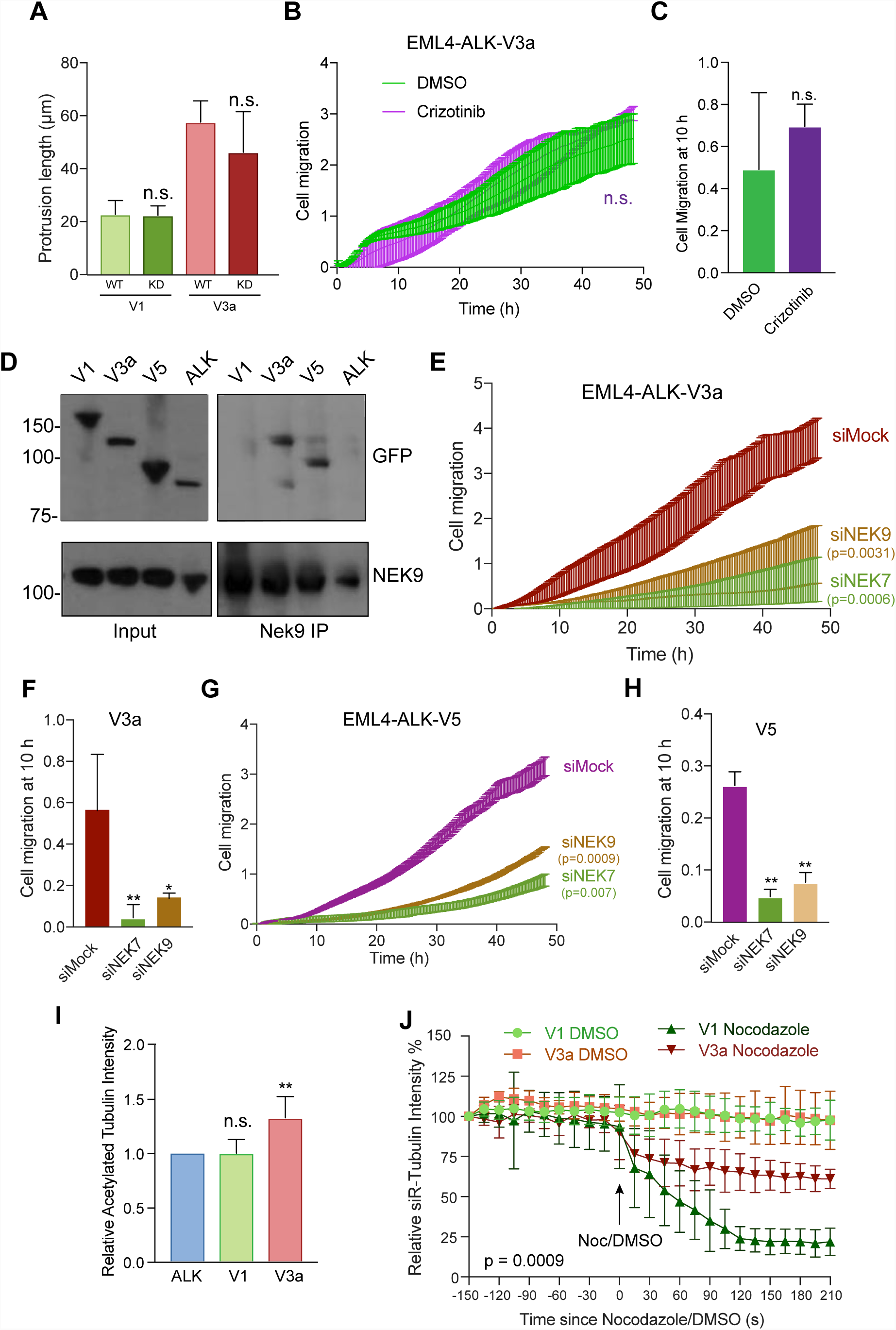
EML4-ALK V3 disturbs cell polarity, stabilises microtubules and binds NEK9. **A**. U2OS cells were transfected with WT and Kinase Dead (KD) versions of YFP-EML4-ALK-V1 and V3a, as indicated, for 72 h prior to analysis by immunofluorescence microscopy. The maximum length of interphase cytoplasmic protrusions for transfected cells was measured. **B**. U2OS:YFP-EML4-ALK-V3a cells were incubated in serum-free medium for 24 h before being treated with DMSO or 100 nM Crizotinib, as indicated, and cell migration analysed in real time as in Fig. 2M. **C**. The histogram shows the cell migration index at 10 h from data in B. **D**. U2OS:YFP-EML4-ALK cells, as indicated, were lysed and lysates (inputs) and immunoprecipitates (IP) prepared with NEK9 antibodies or control IgGs analysed by Western blot with the antibodies indicated. M. wts (kDa) are indicated on the left. **E**. U2OS:YFP-EML4-ALK-V3a cells were mock-, NEK6-or NEK7-depleted for 72 h and cell migration analysed in real time as in Fig. 2M. **F**. The histogram shows the cell migration index at 10 h from data in E. **G**. & **H.** As for E & F but using U2OS:YFP-EML4-ALK-V5 cells. **I**. Cell lysates prepared from U2OS:YFP-EML4-ALK-V1 and V3a cells were Western blotted with acetylated tubulin and GAPDH antibodies and the intensity of acetylated tubulin plotted relative to GAPDH. **J**. U2OS:YFP-EML4-ALK-V1 and V3a were incubated with SiR-Tubulin to visualise microtubules and the SiR-Tubulin intensity measured every 15 s following addition of nocodazole. Control cells were treated with DMSO. Data represent means from 3 separate experiments with 8 wells per condition.

As the EML4-ALK variants all contain the EML4-NTD that in isolation can interact with NEK9, we tested whether these oncogenic fusion variants also bind NEK9. Surprisingly, co-immunoprecipitation experiments showed that endogenous NEK9 interacts with EML4-ALK V3a and V5, but not with V1 or the ALK domain alone (Fig. 7D). The explanation for this might be that, like microtubule binding, interaction with NEK9 is prevented by NTD misfolding or chaperone association of the large variants that possess a partial TAPE domain. Consistent with this and previous studies, real-time transwell migration assays revealed that depletion of NEK9 and NEK7 significantly reduced the migration of cells expressing EML4-ALK V3a or V5 (Fig. 7E-H). As wild-type EML4 can promote microtubule stabilization^36^, we wished to determine whether expression of EML4-ALK V3a caused a similar increase in microtubule stabilisation to that observed in cells expressing activated NEK9. Both Western blot analysis of acetylated tubulin abundance and measurement of the rate of nocodazole-induced microtubule depolymerisation in live cells using the SiR-tubulin probe confirmed that cells expressing EML4-ALK V3a had increased microtubule stability as compared to cells expressed EML4-ALK V1 (Fig. 7I, J; and Supp. Fig. S7B). We therefore conclude that expression of EML4-ALK V3a but not V1 leads to NEK9 and NEK7-dependent alterations in cell morphology and migration, and to enhanced stabilization of microtubules.

### EML4-ALK V3 patient cells exhibit NEK9 and NEK7-dependent alteration of cell morphology and migration

To determine whether the altered cell morphology and migration observed in stable cell lines might contribute to tumour progression in NSCLC patients, we first examined the morphology and migration of established cell lines derived from patients expressing different EML4-ALK variants. H3122 cells are derived from a NSCLC tumour that expresses V1, whereas H2228 are derived from a NSCLC tumour that expresses V3b. Brightfield microscopy revealed that whereas H3122 cells exhibit a cobblestone appearance, H2228 cells have a spindle-like morphology with extended cytoplasmic protrusions (Fig. 8A-C). As observed in the stable cell lines, the protrusions present in H2228 cells were significantly reduced in length upon depletion of either NEK7 or NEK9 (Fig. 8D, E). Real-time transwell migration assays revealed that the H2228 cells exhibited a significantly increased rate of migration as compared to H3122 cells (Fig. 8F, G), while this was reduced by depletion of NEK7 or NEK9 (Fig. 8H, I). Individual cell tracking experiments indicated that depletion of NEK7 or NEK9 reduced the distance and increased the straightness of migration in H2228, but not H3122, cells (Fig 8J, K; and Supp. Fig. S8A-C). Again consistent with data in stable cell lines, the elongated protrusions present in H2228 cells were not affected by treatment with the ALK inhibitor crizotinib, whereas they were reduced upon depolymerisation of microtubules with nocodazole (Fig. 8L, M). Importantly, to confirm that the differences observed in H2228 cells compared to H3122 cells were caused by the different EML4-ALK variant expressed, we first transfected H3122 cells with YFP-EML4-ALK-V3a. This caused a significant increase in the length of interphase cytoplasmic protrusions in H3122 cells (Fig 8N). Second, we depleted EML4-ALK V3 from H2228 cells and found that this led to a significant reduction in cytoplasmic protrusion length (Fig. 8O, P). Together, these data provide persuasive evidence that the different behaviour of these established NSCLC cells results from the different EML4-ALK fusion variant that they express.

**Figure 8.**
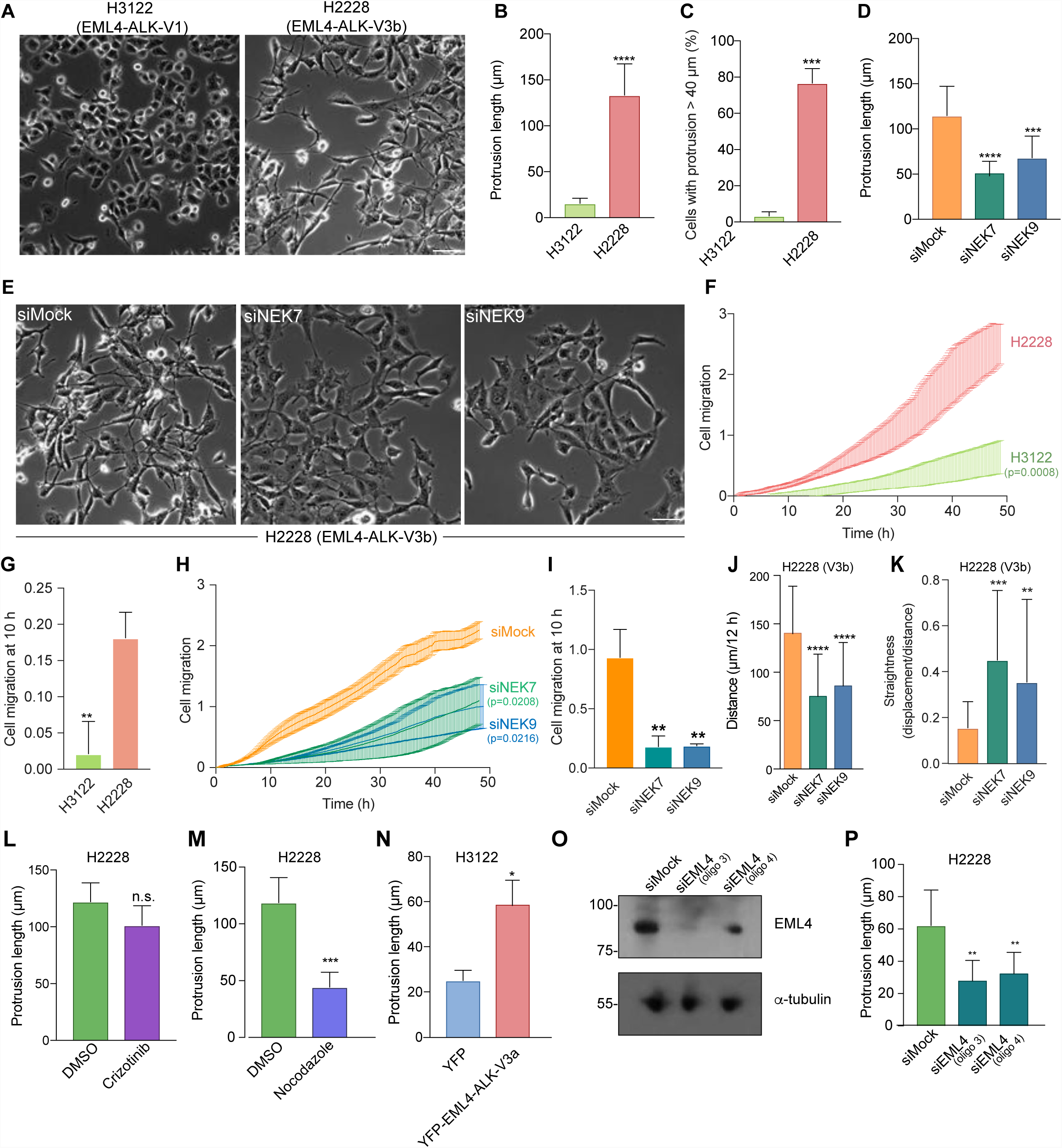
Depletion of NEK7 or NEK9 reduces migration of NSCLC patient cells expressing microtubule-binding EML4-ALK variants. **A**. H3122 and H2228 cells were analysed by phase contrast microscopy. **B**. The maximum length of interphase cytoplasmic protrusions for each cell type was measured. **C**. The % of cells from A and B with protrusions greater than 40 µm is indicated. **D**. H2228 cells were depleted with NEK7 or NEK9 siRNAs for 72 h and the maximum length of cytoplasmic protrusions quantified based on microscopic analysis. **E** Phase contrast images of cells treated as in D are shown. **F**. H3122 and H2228 cell migration was analysed in real time as in Fig. 2M. **G**. Cell index at 10 h for cells in F is shown. **H**. H2228 cells were depleted as indicated for 72 h and migration analysed in real time as in Fig. 2M. **I**. The cell migration index at 10 h for cells in H is shown. **J.** & **K**. H2228 cells that were mock-or NEK9-depleted for 48 h were analysed by individual cell tracking as in Fig. 2I. The mean distance travelled (J) and track straightness (K) is shown **L**. & **M**. H2228 cells were treated with crizotinib (N) or nocodazole (O) and the maximum length of interphase cytoplasmic protrusions measured, control cells were treated with DMSO. **N**. H3122 cells were transfected with either YFP alone or YFP-EML4-ALK V3a and the maximum length of interphase cytoplasmic protrusions for transfected cells measured. **O**. Lysates were prepared from H2228 cells that were mock or EML4-depleted for 72 h and Western blotted as indicated. **P**. Maximum length of interphase cytoplasmic protrusions for cells treated as in O was measured. Scale bars in A, and D, 100 µm.

### Lung cancer patients with EML-ALK V3 exhibit upregulation of NEK9

Results in established cell lines suggest that the more aggressive properties associated with EML4-ALK V3 as compared to V1 may at least in part be caused by a NEK9-dependent pathway that promotes cell migration. This hypothesis would require NEK9 protein to be expressed in EML4-ALK V3 tumours. To test this, we used immunohistochemistry to examine NEK9 expression in a cohort of primary tumours from patients with ALK-positive stage IV advanced lung adenocarcinoma. Remarkably, this revealed that while the majority of tumours expressing V1, V2 (or others) had low levels of NEK9 expression (64%, n=25), the majority of tumours expressing V3 or V5 had medium or high levels of NEK9 expression (91%, n=22; Fig. 9A, B). Intriguingly, patients with medium or high NEK9 expression also exhibited worse overall (p=0.083) and progression-free (p=0.027) survival than patients with low levels of NEK9 expression (Fig. 9C, D). Assessment of clinicopathological features revealed that, although there was no association with gender, age or smoking history, EML4-ALK patients with low NEK9 expression had on average undergone more rounds of previous treatment suggestive of more prolonged and potentially less aggressive disease than those with moderate or high NEK9 expression (Supplementary Tables 1 & 2). Clearly, further work will be required to define the molecular events that underpin the apparent link between elevated NEK9 and both EML4-ALK variant expression and patient survival. However, these findings raise the exciting possibility not only that these NSCLCs could be amenable to treatments that target NEK9 or NEK7, but also that this pathway represents an important mechanism through which variants that bind microtubules accelerate lung cancer progression (Fig. 9E).

**Figure 9.**
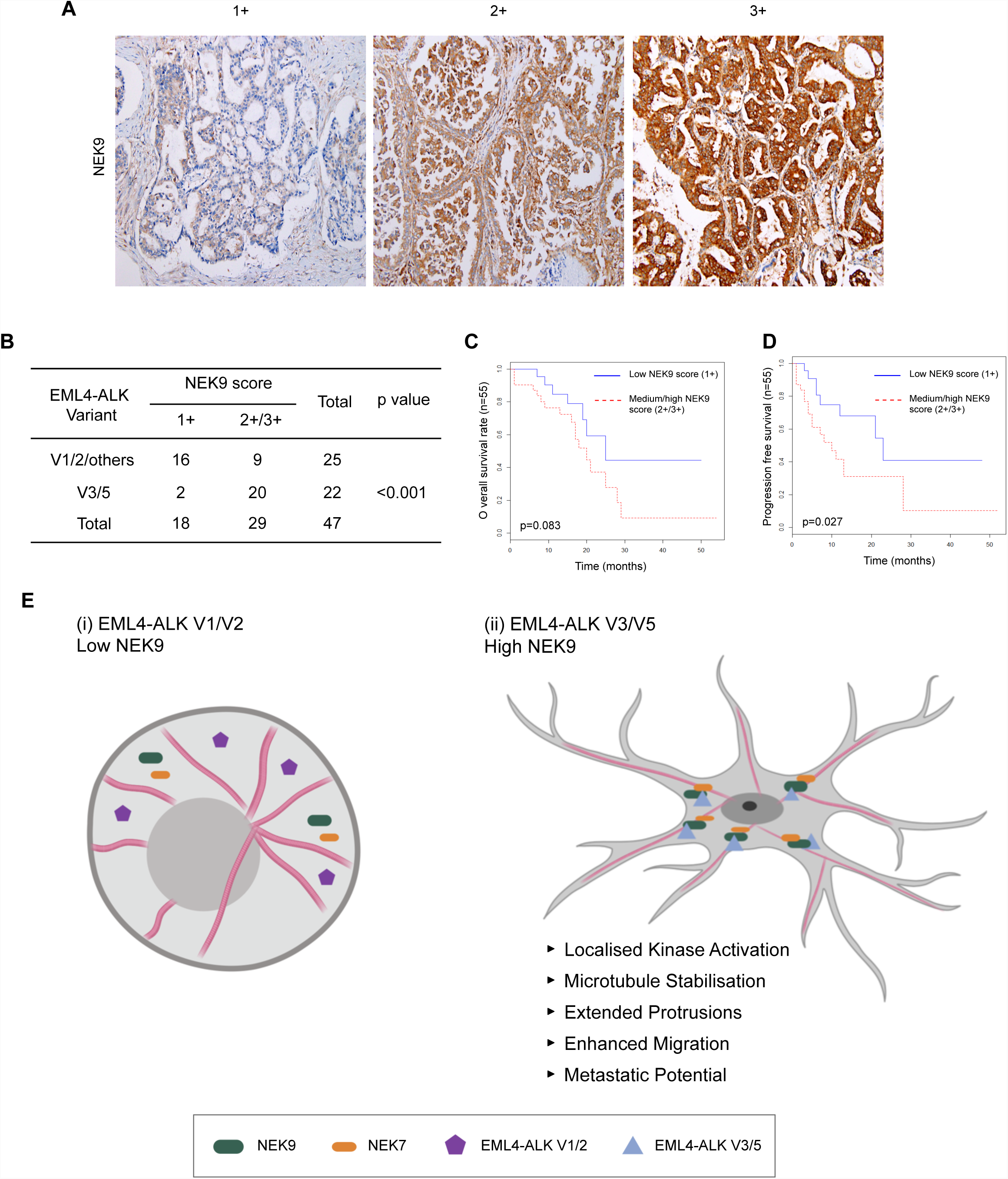
Elevated NEK9 expression correlates with EML4-ALK V3/V5 genotype in NSCLC patient tumours as well as poor overall survival. **A.** Tumour biopsies from NSCLC patients were processed for immunohistochemistry with NEK9 antibodies (brown) and scored as low (1+), medium (2+) or high (3+). Tissue was also stained with haematoxylin to detect nuclei (blue). **B.** Table indicates NEK9 expression as determined as in A with respect to the EML4-ALK variant present. Significance was determined by Fisher Exact Test using R software (n=47). **C.** The Kaplan-Meier plot indicates overall survival of NSCLC patients with EML4-ALK fusion that had low (1+) or medium/high (2+/3+) NEK9 expression (n=32). **D**. As for C but showing progression free survival. **E.** Schematic model showing rationale for stratification of cancer patients for treatment based on EML4-ALK variant. (i) The majority of tumours expressing EML4-ALK V1 or V2 express low levels of NEK9. In these cells, the EML4-ALK protein neither binds NEK9 nor colocalises with microtubules, and cells retain a more rounded morphology. (ii) However, the majority of tumours expressing EML4-ALK V3 or V5 express moderate or high levels of NEK9. In these cells, the EML4-ALK protein binds and recruits NEK9 and NEK7 to microtubules leading to localised kinase activity that promotes microtubule stabilization, extended cytoplasmic protrusions and enhanced migration. This may explain the increased metastatic potential of these tumours.

## DISCUSSION

A better understanding of the processes that drive progression and metastatic dissemination of EML4-ALK tumours is urgently required as this will identify opportunities for development of new therapeutic approaches to treat ALK inhibitor-resistant NSCLC. Here, we show that the short EML4-ALK variants, including the common V3 and less common V5, promote microtubule-dependent changes in cell morphology and enhance cell migration in a manner that could promote dysplastic development and metastasis. EML4-ALK V3 is associated with poor response to ALK inhibitors suggesting additional oncogenic mechanisms that are independent of ALK tyrosine kinase activity^12^. Indeed, we found that these changes induced by EML4-ALK V3 were not dependent on ALK activity. Experimentally, these short variants retain the capacity to bind microtubules. This is in contrast to the longer variants, V1 and V2, that possess destabilizing fragments of the EML4 TAPE domain and rely on the HSP90 chaperone for expression^16^. We speculate that either protein misfolding or chaperone interaction blocks the ability of the longer variants to associate with and regulate microtubules. We also show that the altered cell morphology and increased migration induced by these short EML4-ALK variants are dependent on the NEK9-NEK7 kinase module. Hence, these data not only reveal a novel mechanism through which EML4-ALK variants can promote cancer progression, but also identify a new targeted approach through which patients with these oncogenic fusions could be treated.

NEK9 is required for assembly of the microtubule-based mitotic spindle. This is consistent with the kinase achieving maximal activity in mitosis following phosphorylation by the CDK1 and PLK1 mitotic kinases^24^. These phosphorylation events cause release of an auto-inhibited state that relies on the presence of an internal RCC1-like inhibitory domain^28^. Upon inducible expression of an activated mutant lacking this domain, we observed striking alterations in interphase cell morphology and significantly enhanced cell migration. The mitotic functions of NEK9 are executed in part through phosphorylation of the microtubule nucleation adaptor protein, NEDD1/GCP-WD, and in part through activation of two other members of the NEK family, NEK6 and NEK7^28, 30, 37, 38, 39^. Interestingly, we found that depletion of NEK7, but not NEK6, prevented the altered cell morphology and enhanced migration observed in interphase cells arguing that activated NEK9 promotes these phenotypes through activation of NEK7. Indeed, similar phenotypes were generated by expression of a constitutively active NEK7, but not NEK6, kinase. These findings are consistent with growing evidence that NEK7 has microtubule-associated functions in interphase cells, with endogenous NEK7 localizing to the centrosome and depletion interfering with interphase microtubule dynamics, the centrosome duplication cycle and ciliogenesis^40, 41, 42, 43, 44^. Of particular interest to our findings, a kinase-dependent role was recently described for NEK7 in stimulating dendrite growth in post-mitotic neurons. This resulted from phosphorylation by NEK7 of the kinesin motor protein Eg5, which in turn led to stabilization of microtubules^34^. Supporting a similar mechanism of action in cancer cells, we found that Eg5 was required for the altered cell morphology induced by activated NEK9. As EML4 is highly expressed in neurons, we therefore speculate that modulation of microtubule organization leading to altered cell morphology and migration may represent the normal physiological role for the EML4-NEK9-NEK7 pathway identified here. However, NEK7 can phosphorylate other motor proteins to control microtubule organization, and interacts with additional regulators of both the microtubule and actin networks, offering alternative mechanisms through which these kinases can influence cell movement^45, 46^.

The discovery of EML4 as a binding partner for NEK9 provides new insight into how NEK9 might regulate microtubule dynamics. The EMLs are a family of relatively poorly characterized MAPs^47^. Humans have six EMLs of which EML1, 2, 3 and 4 share a similar organization consisting of an NTD that encompasses a trimerization motif and basic region that binds microtubules, and a C-terminal TAPE domain that binds tubulin heterodimers^15, 16^. The possibility that EMLs may be interacting partners of NEKs was first raised by a proteomic study that identified association of NEK6 not only with NEK7 and NEK9, but also EML2, EML3 and EML4^23^. We identified EML4, as well as tubulin, in a mass spectrometry analysis for NEK9 binding partners, while interaction of endogenous EML4 and NEK9 was confirmed by Western blotting. Co-immunoprecipitation studies revealed that NEK9 can bind to the isolated NTD of EML4, as well as to the EML4-ALK V3 and V5 proteins. Interestingly, it did not bind to EML4-ALK V1 despite this variant encoding the complete NTD. Again this suggests that misfolding or binding of chaperones prevents interaction with NEK9. Further studies revealed that the EML4 NTD was capable of recruiting both NEK9 and NEK7 to microtubules. Firstly, this suggests that binding of the NTD to microtubules and NEK9 are not mutually exclusive. Secondly, it allows us to propose that EML4 recruits NEK9 to microtubules and that, in turn, NEK9 recruits NEK7. Assuming that the morphological consequences that we observe are dependent upon phosphorylation of microtubule-associated substrates downstream of NEK7, this also explains why NEK9 is required even in the presence of an activated NEK7 mutant. However, although we have shown that expression of activated NEK9 and EML4-ALK V3 leads to stabilization of microtubules, further work is required to establish the detailed mechanisms through which this pathway alters microtubule dynamics to change cell shape, polarity and migration.

Of major clinical significance is the fact that EML4-ALK fusion proteins act as oncogenic drivers in ∼5% of NSCLC patients^4^. Unscheduled activation of the ALK tyrosine kinase is primarily responsible for driving tumorigenesis as demonstrated by the fact that ALK inhibitors block transformation. ALK inhibitors have revolutionized survival outcomes in EML4-ALK NSCLC, however resistance is inevitable emerging through point mutations in the catalytic domain as well as through activation of bypass signalling pathways^48, 49^. Alternative targeted therapies would therefore be highly attractive and, in the case of the longer variants, HSP90 inhibitors could provide an option. The longer variants are unstable proteins as a result of expressing an incomplete fragment of the highly structured TAPE domain. In the absence of the HSP90 chaperone they are rapidly degraded by the proteasome leading to loss of downstream proliferative signalling^16^. HSP90 inhibitors have been tested in EML4-ALK-positive patients although to date clinical trials have yet to prove their efficacy^19, 50^. The short variants though are not dependent on HSP90 for their expression and so HSP90 inhibitor treatment is unlikely to be effective. Moreover, EML4-ALK V3 tumours are insensitive to ALK inhibitors in the first place emphasizing the need for alternative treatments for these particular patients^12^. Demonstration that the downstream consequences on cell morphology and migration induced by these short variants is dependent on NEK9 and NEK7, but not ALK activity, raises the tantalizing prospect that inhibitors of this pathway would be beneficial in EML4-ALK V3 cancers where metastasis and ALK inhibitor resistance underpin lethality^22^. Although we have no evidence that inhibiting this pathway promotes cytotoxicity, the ability to block migration and invasion with so-called ‘migrastatics’ offers huge potential benefits in slowing disease progression in patients with solid cancers^51^.

The significant correlation in patient samples between elevated NEK9 expression and EML4-ALK V3 is particularly intriguing as it implicates this pathway in the high metastatic potential of these tumours^22, 52^. However, although the association of elevated NEK9 with V3 and V5 was significant, expression does not necessarily reflect activation and it will be important in the future to assess the catalytic activity of NEK9 and NEK7 in these tumours. Moreover, we cannot rule out that tumour cells with this variant exert positive selection for high expression of NEK9 either due to its role as a downstream target or as a result of their increased aggressiveness. Nevertheless, the reduced progression-free and overall survival of patients with increased NEK9 expression potentially explains the poor response of these patients to ALK-based therapies, despite the fact that the different EML4-ALK variants bind ALK inhibitors with similar affinity^17^. This also adds further evidence that NEK9 may be an important prognostic factor for EML4-ALK lung cancers and supports the hypothesis that targeting NEK9 or NEK7 could be beneficial in patients with EML4-ALK V3.

Finally, it is interesting to note that EML proteins are involved in other oncogenic fusions, including the EML1-ABL1 fusion found in T-cell acute lymphoblastic leukaemia (ALL)^53^. Whether these also localise to microtubules and alter cell morphology and migration in a manner that is dependent on the microtubule network and the NEK9-NEK7 kinase module will be important to examine. Standard treatments for many cancers, including NSCLC and ALL, often involve microtubule poisons, such as vinca alkaloids or taxanes. Hence, it will also be worthwhile to explore the possibility that EML4-ALK variant status affects response to these drugs, and test whether there could be patient benefit in combining microtubule poisons with targeted agents against ALK, HSP90 or indeed NEK9 or NEK7.

## MATERIALS & METHODS

### Plasmid construction and mutagenesis

Full-length EML4 cDNA was isolated by PCR from human cDNA (Clontech) and subcloned into a version of pcDNA3 or pcDNA3.1-hygro (Invitrogen) providing N-terminal YFP-or Flag-tags. YFP-tagged EML4-ALK variants and fragments of EML4, and Flag-tagged NEK6 and NEK7 constructs were generated as previously described^15, 16, 54^. A full-length Myc tagged cDNA expressing human NEK9 (NM_001329237.1) was subcloned into the pVLX-Tight-Puro vector (Clontech). The RCC1 domain was deleted using the Quickchange^®^ II XL Site-Directed Mutagenesis Kit according to manufacturer’s instructions (Stratagene) using the primer sequences, 5’-gctgtagtaacatcacgaaccagtatccgttccaatagcagtggcttatcc and 5’-ggataagccactgctattggaacggatactggttcgtgatgttactacagc. Point mutations in the NEK7, NEK6 and ALK catalytic domains were also introduced using the Quickchange^®^ II XL Site-Directed Mutagenesis Kit. Constructs were confirmed by DNA sequencing (University of Leicester).

### Cell culture, drug treatments and transfection

HeLa, U2OS and derived stable cell lines were grown in Dulbecco’s modified Eagle’s medium (DMEM) with GlutaMAX™-I (Invitrogen) supplemented with 10% heat-inactivated fetal bovine serum (FBS), 100 U/ml penicillin, and 100 µg/ml streptomycin, at 37°C in a 5% CO_2_ atmosphere. Doxycycline-inducible NEK9 stable U2OS cell lines were generated through lentiviral transduction of parental U2OS cells stably carrying the pVLX-Tet-On-Advance vector (Clontech, PT3990-5). Briefly, NEK9 lentiviral particles for each construct were harvested from transfected 293T packaging cells using the Lenti-X(tm) HTX Packaging System (Clontech), transduced into U2OS parental cells and selected in fresh media supplemented with 2 mg/ml of G418 and 1 µg/ml puromycin over several days. To maintain expression of constructs, culture media was supplemented with 1 ug/ml of G418 and 800 ng/ml of puromycin. For induction of constructs, doxycycline was added to growth media at a final concentration of 1 µg/ml for 72 h, unless otherwise stated. Doxycycline-inducible HeLa:YFP-NEK7 cell lines were generated via co-transfection of parental HeLa cells which stably express the lacZ-Zeocin™ fusion gene and Tet repressor from pFRT/lacZeo and pcDNA6/TR plasmids, respectively, with the appropriate pcDNA5/FRT/TO-YFP-NEK7 vector and the Flp recombinase expression vector, pOG44. After 48 h cells were incubated in selective media containing 200 µg/ml hygromycin B and resistant clones selected and expanded. To maintain expression in stable cell lines, media was supplemented with 200 µm/ml hygromycin B (Invitrogen). For induction of NEK7 constructs, doxycycline was added at a final concentration of 1 µg/ml for 72 h. Constitutively expressing EML4 and EML4-ALK cell lines were generated by transfecting the relevant construct into U2OS cells. After 48 h cells were incubated in selective media containing 100 µg/ml hygromycin B (Invitrogen). To maintain expression in stable cell lines, media was supplemented with 100 µg/ml hygromycin. Where indicated, cells were treated with 200 ng/ml nocodazole (Sigma), 20 nM vinorelbine (Sigma), 100 nM STLC (Tocris) or 100 nM Crizotinib (Selleck). Transient transfections were performed with Lipofectamine 2000 (Invitrogen) according to manufacturer’s instructions. For flow cytometry, cells were fixed in ice-cold 70% ethanol before DNA was labelled with 5 *μ*g/ml propidium iodide. Analysis was performed using a FACSCanto II instrument and FACSDiva software (BD Biosciences).

### Preparation of cell extracts, SDS-PAGE and Western blotting

Cells were lysed in RIPA lysis buffer (50 mM Tris-HCl pH 8, 150 mM NaCl, 1% v/v Nonidet P-40, 0.1% w/v SDS, 0.5% w/v sodium deoxycholate, 5 mM NaF, 5 mM β-glycerophosphate, 30 µg/ml RNase, 30 µg/ml DNase I, 1x Protease Inhibitor Cocktail, 1 mM PMSF) prior to analysis by SDS-PAGE and Western blotting. Primary antibodies were GFP (0.2 µg/ml, Abcam), α-tubulin (0.1 µg/ml, Sigma), myc (1:1000, Cell Signalling Technologies), NEK9 (0.8 µg/ml, Santa Cruz Biotechnology), EML4 (0.2 µg/ml, Bethyl Laboratories), Flag (0.5 µm/ml, Sigma), acdetylated tubulin (0.5 µg/ml, Sigma) ALK (1:2000, Cell Signaling technology), pALK (1:1000, Cell Signaling Technology) and GAPDH (1 µg/ml, Cell Signalling Technology). Secondary antibodies used were anti-rabbit, anti-mouse or anti-goat horseradish peroxidase (HRP)-labelled IgGs (1:1000; Sigma). Western blots were detected using enhanced chemiluminescence (Pierce).

### RNAi

Cells at 30–40% confluency were cultured in Opti-MEM Reduced Serum Medium and transfected with 50 nM siRNA duplexes using Oligofectamine (Invitrogen) according to the manufacturer’s instructions. 72 hours after transfection, cells were either fixed for immunocytochemistry or prepared for Western blot. siRNA oligos were directed against NEK9, AM51334-1113 and -1115 silencer select duplexes (Ambion) or EML4, HSS120688 and Hss178451 On-Target Plus duplexes (Dharmacon); siRNA duplexes for NEK6 and NEK7 were as previously described^54, 20^.

### Immunoprecipitation, kinase assay and mass spectrometry

Cells were harvested by incubation with 1x PBS+0.5 mM EDTA and pelleted by centrifugation prior to being lysed in NEB lysis buffer^55^. Lysates were immunoprecipitated using antibodies against NEK9 (0.8 µg/ml, goat, Santa Cruz Biotechnologies), EML4 (0.4 µg/ml, rabbit, Bethyl Laboratories), myc (1:1000, mouse, Cell Signaling Technologies) or GFP (0.2 µg/ml, rabbit, Abcam). Proteins that co-precipitated with myc-NEK9 were excised from gels and subjected to in-gel tryptic digestion prior to LC-MS/MS using an RSLCnano HPLC system (Dionex, UK) and an LTQ-Orbitrap-Velos mass spectrometer (Thermo Scientific). The raw data file obtained from each LC-MS/MS acquisition was processed using Proteome Discoverer (version 1.4, Thermo Scientific), searching each file in turn using Mascot (version 2.2.04, Matrix Science Ltd.) against the human reference proteome downloaded from UniProtKB (Proteome ID: UP000005640). The peptide tolerance was set to 5 ppm and the MS/MS tolerance set to 0.6 Da. The output from Proteome Discoverer was further processed using Scaffold Q+S (version 4.0.5, Proteome Software). Upon import, the data was searched using X!Tandem (The Global Proteome Machine Organization). PeptideProphet and ProteinProphet (Institute for Systems Biology) probability thresholds of 95% were calculated from the decoy searches and Scaffold used to calculate an improved 95% peptide and protein probability threshold based on the two search algorithms.

### Fixed and time-lapse microscopy

Cells grown on acid-etched coverslips were fixed and permeabilised by incubation in ice-cold methanol at −20°C for a minimum of 20 min. Cells were blocked with PBS supplemented with 3% BSA and 0.2% Triton X-100 prior to incubation with the appropriate primary antibody diluted in PBS supplemented with 3% BSA and 0.2% Triton X-100. Primary antibodies used were against GFP (0.5 µg/ml, rabbit; Abcam); α-tubulin (0.1 µg/ml, mouse, Sigma), myc (1:1000, mouse, Cell Signalling Technologies), Flag (0.5 µg/ml, mouse, Sigma-Aldrich), NEK9 (0.8 µg/ml, goat, Cell Signalling Technology), GM130 (0.4 µg/ml, rabbit, Abcam), γ-tubulin (0.2 µg/ml, Sigma). Secondary antibodies were Alexa Fluor 488, 594 and 647 donkey anti-rabbit, donkey anti-mouse and donkey anti-goat IgGs (1 µg/ml; Invitrogen). DNA was stained with 0.8 µg/ml Hoechst 33258. For actin visualisation, cells were fixed using 3.7% formaldehyde and stained using phalloidin-TRITC (1:200). Imaging was performed on a Leica TCS SP5 confocal microscope equipped with an inverted microscope (DMI6000 B; Leica) using a 63x oil objective (numerical aperture, 1.4). Z stacks comprising 30–50 0.3-µm sections were acquired using LAS-AF software (Leica), and deconvolution of 3D image stacks performed using Huygens software (Scientific Volume Imaging). For super-resolution radial fluctuations (SRRF) microscopy a VisiTech infinity 3 confocal microscope fitted with Hamamatsu C11440 -22CU Flash 4.0 V2 sCMOS camera and a Plan Apo 100 x objective (NA 1.47) was used. 100 images from the same slice were captured per channel and processed using the nanoJ-SRRF plugin in Fiji. Immunohistochemistry with NEK9 antibodies (Abcam, ab138488; 1:500) was performed on NSCLC patients with EML4-ALK fusions.

Phase contrast and time-lapse microscopy was carried out using a Nikon eclipse Ti inverted microscope using a Plan Fluor 10x DIC objective (NA 0.3) or a Plan Fluor 40x objective (NA 1.3). Images were captured using an Andor iXonEM+ EMCCD DU 885 camera and NIS elements software (Nikon). For time-lapse imaging, cells were cultured in 6-well dishes and maintained on the stage at 37°C in an atmosphere supplemented with 5% CO_2_ using a microscope temperature control system (Life Imaging Services) with images acquired every 15 min for ≥24 h. Videos were prepared using ImageJ (National Institutes of Health). Cell protrusions were measured using Fiji. The length of protrusion was defined as the distance from the edge of the nucleus of the cell to the furthest point of the plasma membrane.

### Individual Cell Tracking Assays

To allow automated detection of individual cells for tracking analysis cells were transfected with YFP alone and subjected to time lapse imaging as appropriate. Individual cell tracking was then analysed using Imaris (Bitplane). Fluorescent cells were identified using the Imaris spots function and subsequently the Imaris TrackLineage algorithm was used to identify movements or tracks of these cells through the entire time period and thus provide numerical values for parameters such as speed and distance.

### Live-cell microtubule stability assays

For live cell microtubule stability assays cells were grown in µ-well 8 well chamber slides (ibidi). Cells were incubated with 25 nM SiR-Tubulin (Cytoskeleton Inc.) for 4 h prior to imaging, 10 images were captured prior to addition of nocodazole and the drug was then added and a further 15 images were captured. Z stacks comprising of approximately 10 0.5 µm sections were captured every 30 s for 7 mins. Images were cropped to single cells and deconvolved prior to analysis in Matlab.

### Cell migration assays

For scratch-wound assays, cells were grown to 95% confluency in 6 well dishes before a pipette tip was used to scrape a 0.5 - 1 µm line across the width of the well. Cells were washed 3-5 x in pre-warmed media and either imaged by time-lapse microscopy or incubated at 37°C for 6 h before being processed for immunofluorescence microscopy. For Golgi and centrosome orientation measurements, the cell was divided into 4 quadrants (Q), with Q1 oriented towards the wound. Cells were scored as having Golgi or centrosomes oriented towards the wound if the majority of the Golgi or both centrosomes, respectively, fell into Q1. Cell migration in real time was analysed using the xCELLigence Real-Time Cell Analyzer (RTCA) DC equipment (AECA Biosciences) and CIM-16 plates, a 16 well system where each well is composed of upper and lower chambers separated by an 8 µm microporous membrane. Cells were grown in serum free medium for 24 h before being seeded in serum free medium into the upper chamber; complete medium containing 10% FBS was used as a chemo-attractant in the lower chamber of the plate and migration determined as the relative impedance change (cell index) across microelectronic sensors integrated into the membrane. Measurements were taken every 15 mins for 48 h.

### Statistical analysis

All quantitative data represent means and SD of at least three independent experiments. Statistical analyses were performed using a one-tailed unpaired Student’s *t* test assuming unequal variance, a one-way analysis of variance followed by post hoc testing or a χ^2^ test, as appropriate. *p*-values represent *, *p*< 0.05; **, *p*< 0.01; ***, *p*< 0.001. n.s., non-significant.

## ACKNOWLEDGMENTS

We are very grateful to Patrick Meraldi and his colleagues (University of Geneva) for introducing us to the kinetic SiR-tubulin microtubule depolymerization assay. We acknowledge the University of Leicester Core Biotechnology Services for support with cloning, mass spectrometry, DNA sequencing and microscopy. This work was supported by grants to A.M.F. from Worldwide Cancer Research (13-0042 and 16-0119), the Wellcome Trust (082828 and 097828) and Cancer Research UK (C1362/A180081); to R.B. from Cancer Research UK (C24461/A23302); to R.B. and J.C. from MRC-KHIDI (MC_PC_17103) U.K. and MRC-KHIDI (HI17C1975) Republic of Korea; to S.J.C. from Cancer Research UK (Senior Cancer Research Fellowship A12102); and to G.B. from Weston Park Hospital Cancer Charity (Large Project Grant CA164).

## AUTHOR CONTRIBUTIONS

L.O’R., G.B, R.A., C.G.W., H.J.J. and E.L.R. undertook the experiments and provided comments on the manuscript. M.R. designed and generated the YFP-EML4 and YFP-EML4-ALK constructs; P.A.J.M., S.J.C., D.A.F., J.C. and R.B. contributed to study design and provided comments on the manuscript. A.M.F. designed and supervised the project and wrote the manuscript.

