## Supplementary Material for "EML4-ALK V3 drives cell migration through NEK9 and NEK7 kinases in non-small-cell lung cancer"

### SUPPLEMENTARY FIGURE LEGENDS

**Figure S1. Characterisation of doxycycline-inducible U2OS:myc-NEK9 constructs**

**A**. U2OS:myc-NEK9-WT, ΔRCC1 or K81M cells, or cells containing empty vector, were either uninduced (- dox) or induced (+ dox) for 72 h before being processed for immunofluorescence microscopy with myc (red) antibodies. **B**. Cells were treated as in A and analysed by immunofluorescence microscopy with myc (green) and α-tubulin (red) antibodies. In A & B, DNA was stained with Hoechst 33258 (blue); scale bars, 20 µm. **C.** U2OS:myc-NEK9-WT, ΔRCC1 or K81M cells were induced for 72 h before being fixed, stained with propidium iodide and analysed by flow cytometry. The percentage of cells in G1, S and G2/M are indicated. **D**. U2OS:myc-NEK9-ΔRCC1 cells were treated for 48 hours +/- doxycycline before incubation with SiR-Tubulin to visualise microtubules. SiR-Tubulin intensity was then measured every 15 s following addition of nocodazole. Stills from movies are shown. Scale bar, 10 µm.

**Figure S2. Depletion of Nek6 and Nek7 in Nek9-ΔRCC1 cell lines**

**A.** U2OS:myc-NEK9-ΔRCC1 cells were mock-depleted or depleted with NEK6 or NEK7 siRNAs for 24 h prior to induction with doxycycline for a further 48 h. Cell lysates were prepared and analysed by Western blot with myc, NEK6, NEK7 and α-tubulin antibodies. **B**. U2OS:myc-NEK9-ΔRCC1 cells were mock-depleted, or depleted with siNEK6.1 or siNEK7.1 and processed for immunofluorescence microscopy with myc (red) and α-tubulin (green) antibodies. DNA was stained with Hoechst 33258. Scale bar, 20 µm. **C**. U2OS:myc-NEK9-ΔRCC1 cells were treated as in A and analysed using the live cell transwell migration assay. Data represent means from 3 separate experiments. **D**. Histogram shows the cell migration index at 10 h based on data in C.

**Figure S3. Characterisation of doxycycline-inducible HeLa:YFP-NEK7 constructs**

Lysates prepared from HeLa stable cell lines induced to express wild-type (WT), catalytically-inactive (D179N) or constitutively-active (Y97A) YFP-NEK7 with doxycycline for 0, 24 and 48 h were analysed by Western blot with GFP and α-tubulin antibodies. M. wts (kDa) are indicated on the left.

**Figure S4. Identification of EML4 as a NEK9 binding partner by mass spectrometry**

**A**. Anti-myc immunoprecipitates prepared from U2OS:myc-NEK9-WT cells induced with doxycycline for 48 hours were analysed by SDS-PAGE and mass spectrometry. Eight peptides representing EML4 were identified in the immunoprecipitates from myc-NEK9 expressing cells, whereas no EML4 peptides were identified in immunoprecipitates from parental U2OS cells. **B**. The table shows peptides identified from human EML4 by LC-

MS/MS. **C**. MS/MS spectrum of peptide WFVLDAETR; y-ion fragments highlighted in blue, b-ion fragments in red. Inset shows full fragmentation table.

**Figure S5. Characterization of U2OS:YFP and U2OS:YFP-EML4 stable cell lines**

**A**. Lysates prepared from U2OS:YFP and U2OS:YFP-EML4 cell lines were Western blotted with the antibodies indicated. **B**. Immunofluorescence microscopy images of U2OS:YFP and U2OS:YFP-EML4 cell lines stained with GFP (green) and $\alpha$-tubulin (red) antibodies. DNA (blue) is stained with Hoechst 33258. Scale bar, 20 $\mu$m. **C**. Representative intensity profiles along a single line-scan showing co-localization of YFP or YFP-EML4 with microtubules. *R* values show the mean Pearson's correlation coefficient from 10 cells. **D**. Phase contrast microscopy of U2OS:YFP and U2OS:YFP-EML4 cell lines. Scale bar, 100 µm. **E**. Lysates prepared from U2OS:myc-NEK9-ΔRCC1 cells depleted for EML4 with two siRNAs or mock-depleted for 24 h prior to induction with doxycycline for a further 48 h were analysed by Western blot for endogenous EML4 and actin. **F**. Lysates prepared from U2OS:YFP and U2OS:YFP-EML4 cells, which had been depleted for NEK7 or NEK9 or mock depleted for 72 h, were Western blotted for NEK7, NEK9, GFP or $\alpha$-tubulin. **G**. U2OS:YFP or U2OS:YFP-EML4 cell lines were depleted of NEK7 or NEK9, as indicated, before analysis by phase contrast microscopy; scale bar, 100 µm. M. wts (kDa) are indicated on the left in A, E and F.

**Figure S6. EML4-NTD recruits NEK9 and NEK7 to microtubules**

**A.** U2OS cells were transfected with YFP-EML4 N-terminal domain (NTD) or TAPE domain (TAPE), as indicated, for 24 h before being processed for immunofluorescence microscopy with antibodies against GFP (green) and NEK9 (red). **B**. HeLa:YFP-NEK7 cells were induced for 48 h with doxycycline before being mock transfected or transfected with Flag-EML4-NTD or TAPE for a further 24 h. Cells were then processed for immunofluorescence microscopy with antibodies against Flag (green) and GFP (red). Merge images include DNA stained with Hoechst 33258 (blue). Scale bars, 10 µm.

**Figure S7. Inhibition of ERK phosphorylation by crizotinib and stabilization of microtubules in EML4-ALK V3 cells**

**A**. Cell lysates prepared from U2OS:YFP-EML4-ALK-V3a cells treated with DMSO or Crizotinib at the concentrations indicated for 8 h were analysed by Western blot with ERK, pERK and $\alpha$-tubulin antibodies. **B**. U2OS:YFP-EML4-ALK V1 or V3a cells were incubated with SiR-Tubulin to visualise microtubules before SiR-Tubulin intensity was measured every 15 s following addition of DMSO or nocodazole. Stills from movies are shown. Scale bars, 10 µm.

**Figure S8. Validation of siRNA depletion of NEK7 and NEK9 in EML4-ALK NSCLC cells**

**A**. H3122 or H2228 cell lysates depleted for NEK6, NEK7 or NEK9 as indicated were analysed by Western blot with NEK7, NEK9 and $\alpha$-tubulin antibodies. M. wts (kDa) in A & B are indicated on the left. **B**. & **C**. H3122 cells that were mock, NEK7 or NEK9-depleted for 48 h were analysed by individual cell tracking. The mean distance travelled (C) and track straightness (D) is shown.

**Supplementary Table S1. Clinicopathological characteristics of EML4-ALK NSCLC patients**

182 ALK-positive patients with advanced NSCLC at Asan Medical Center (Seoul, Korea) were collected between June 2011 and August 2015. Of those, 113 patients were treated with the ALK inhibitor and had an Eastern Cooperative Oncology Group (ECOG) performance status between 0 and 3. Among 93 enrolled patients who were tissue-available and approved by the institutional review board, 38 were excluded because of poor quality of insufficient tissue samples and follow-up loss. For the analysis of the remaining 55 patients, medical records were reviewed to extract clinicopathological data including sex, age, smoking status, therapeutic agents, and survival. Of those, ALK subtyping was performed in 32 samples due to 23 cases with low quality of nucleic acid. All statistical analyses were carried out in the R software (version 3.3.3, the R Foundation for Statistical Computing, Vienna, Austria).

**Supplementary Table S2. Clinicopathological characteristics of EML4-ALK NSCLC patients according to NEK9 expression**

Based on the patients described in Supplementary Table S1, no clinicopathological characteristics exhibited a significant difference in relation to NEK9 expression. However, the number treated with previous chemotherapy was close to the borderline for significance ($p$=0.071) in that the proportion of patients treated with first line ALK inhibitor therapy was larger in the high Nek9 expression (score 2+/3+) group than the low expression (score 1+) group. This means that the low expression group included more intensively treated patients suggestive of more advanced NSCLC. This factor may affect the survival difference in the two groups treated with the ALK inhibitor.

**Supplementary Movie S1. Time-lapse imaging of cells expressing NEK9-ΔRCC1**

U2OS:myc-NEK9-ΔRCC1 cells were treated with doxycycline and imaged by time-lapse phase contrast microscopy. Images were acquired every 15 minutes for 24 h.

**Supplementary Movie S2. Time lapse imaging of control cells**

U2OS:myc-NEK9-ΔRCC1 cells were treated with DMSO and imaged by time-lapse phase contrast microscopy. Images were acquired every 15 minutes for 24 h.

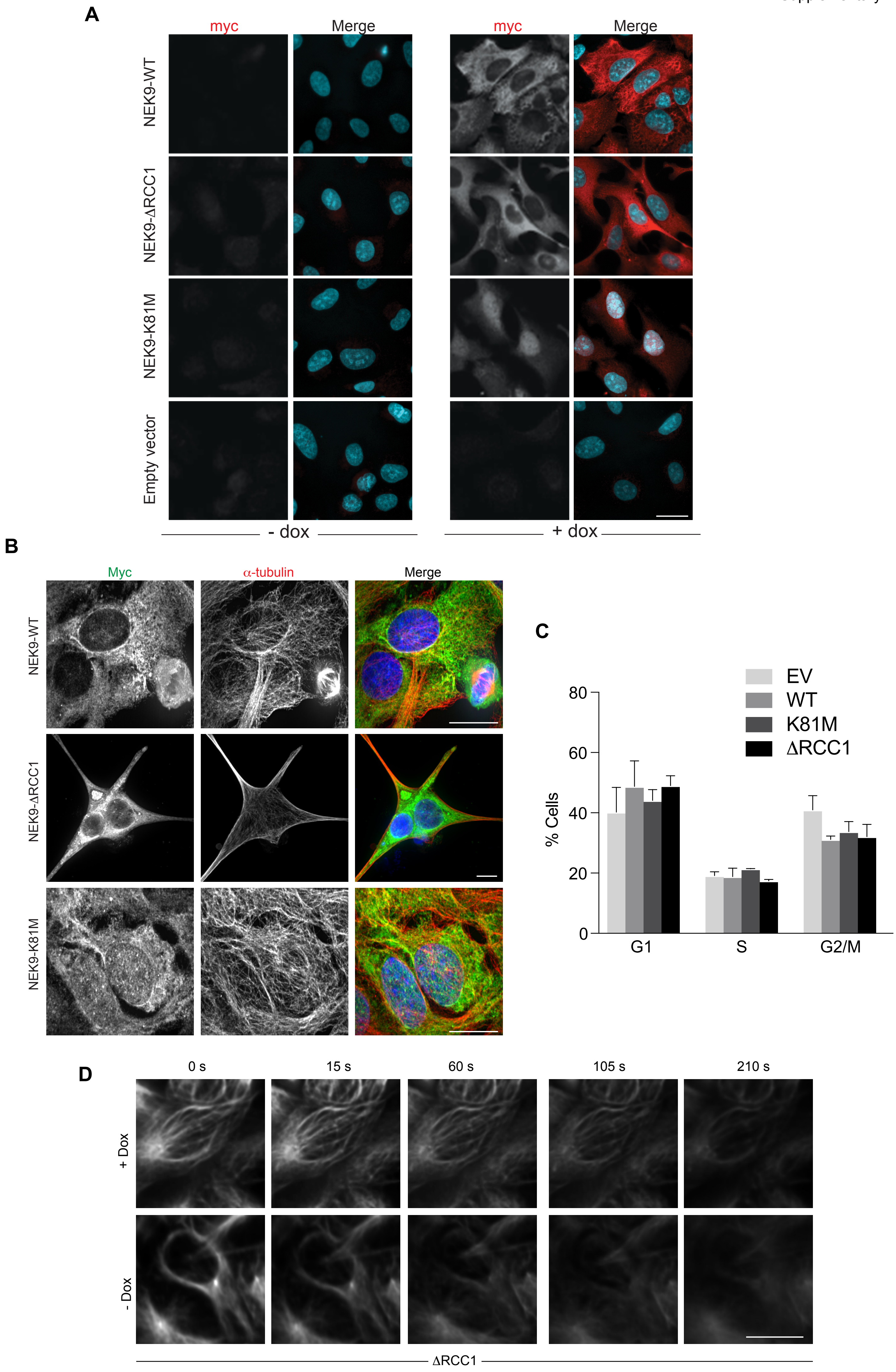
A
myc
Merge
myc
Merge
NEK9-WT
NEK9-ΔRCC1
NEK9-K81M
Empty vector
- dox
+ dox
B
Myc
α-tubulin
Merge
NEK9-WT
NEK9-ΔRCC1
NEK9-K81M
C
EV
WT
K81M
ΔRCC1
80
60
40
20
0
% Cells
G1
S
G2/M
D
0 s
15 s
60 s
105 s
210 s
+ Dox
- Dox
ΔRCC1

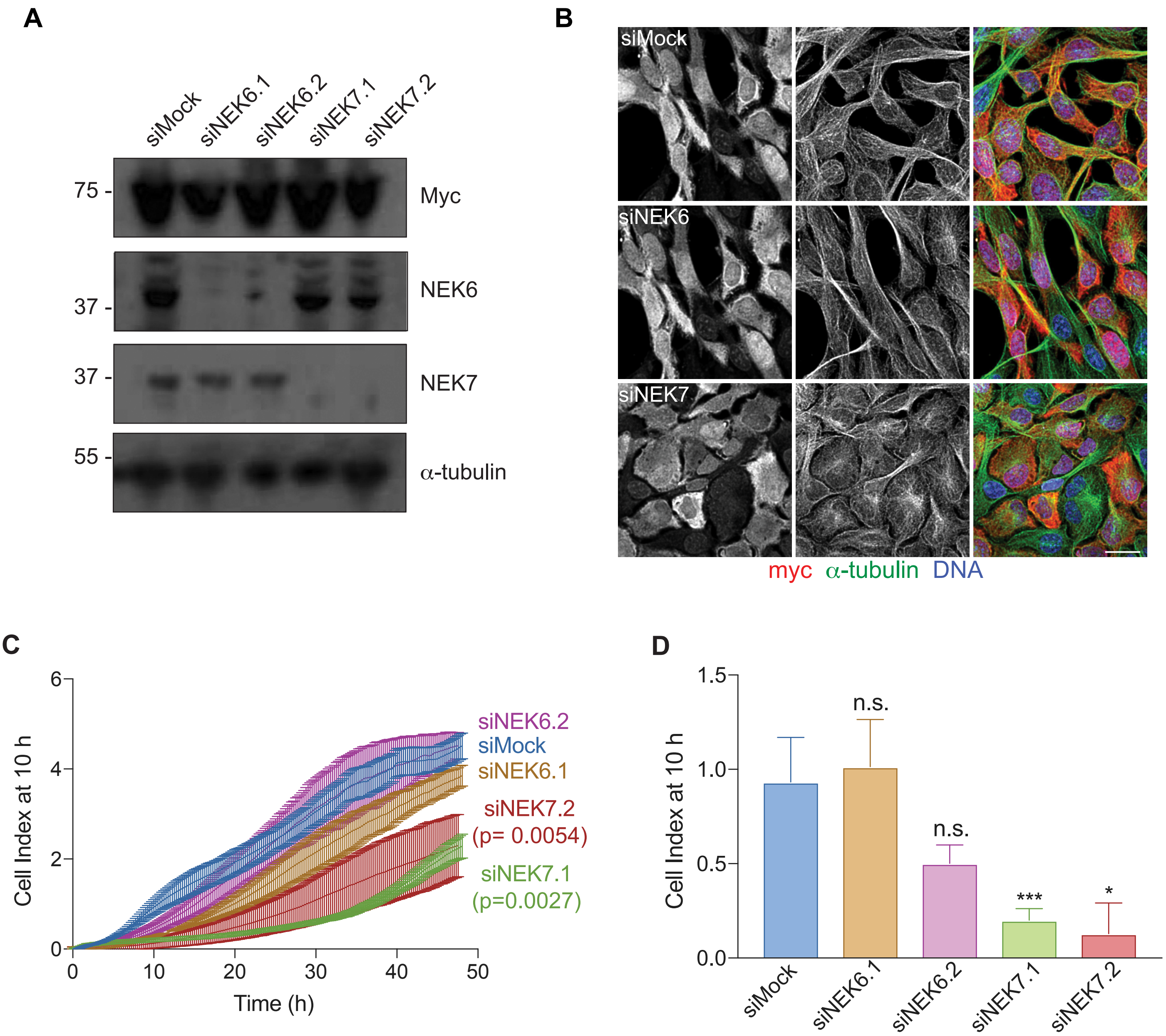
A
siMock
siNEK6.1
siNEK6.2
siNEK7.1
siNEK7.2
75 -
Myc
NEK6
37 -
37 -
NEK7
55 -
α-tubulin
B
siMock
siNEK6
siNEK7
myc α-tubulin DNA
C
Cell Index at 10 h
Time (h)
siNEK6.2
siMock
siNEK6.1
siNEK7.2
(p= 0.0054)
siNEK7.1
(p=0.0027)
D
Cell Index at 10 h
n.s.
n.s.
***
*
siMock
siNEK6.1
siNEK6.2
siNEK7.1
siNEK7.2

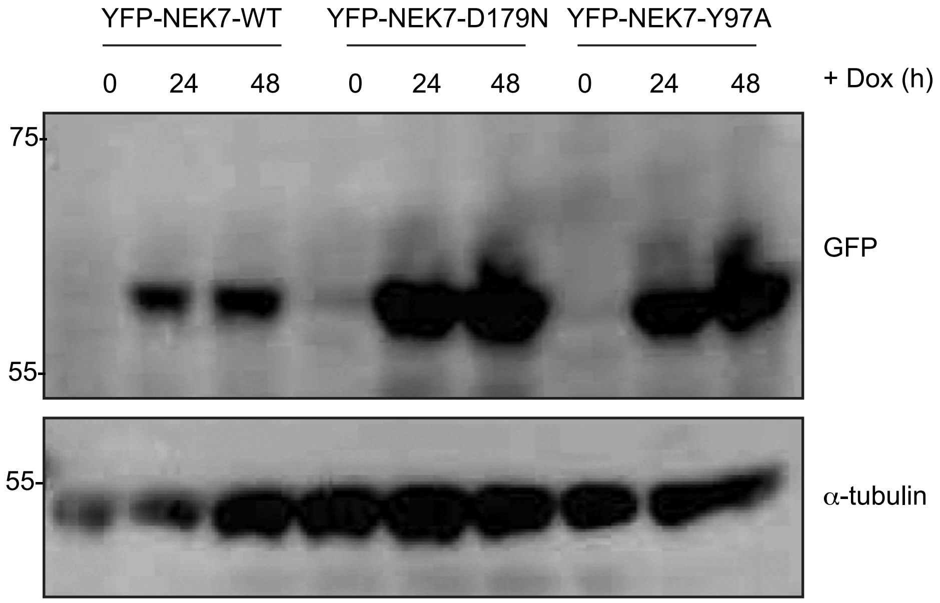
YFP-NEK7-WT
YFP-NEK7-D179N
YFP-NEK7-Y97A
0 24 48 0 24 48 0 24 48
+ Dox (h)
75
55
GFP
55
α-tubulin

**A**

Probablilty legend:

| Probablilty legend: |
|---|
| Over 95 % |
| 80-94 % |
| 50-79 % |
| 20-49 % |
| 0-19 % |

| UL00825GB | | | | Control | Nek9 |
|---|---|---|---|---|---|
| # | Identified Proteins (3) | Accession Numbe | Molecular Weight | UL00825GB - 02 - Con | UL00825GB - 01 - Nek9 |
| 1 | Serine/threonine-protein kinase Nek9 OS=Homo sapiens GN=NEK9 PE=1 SV=2 | Q8TD19 | 107 kDa | ! | 29 |
| 2 | Echinoderm microtubule-associated protein-like 4 OS=Homo sapiens GN=EML4 PE=1 SV=3 | Q9HC35 | 109 kDa | 0 | 8 |

**B**

| Sequence | Scaffold Probability | Mascot Ion score | Mascot Identity score | X! Tandem | Observed | Actual Mass | Charge | Delta Da | Delta PPM |
|---|---|---|---|---|---|---|---|---|---|
| (K)DVIINQEGEYIK(M) | 100% | 29.2 | 22.4 | 3.72 | 710.8697 | 1419.7248 | 2 | 0.0001 | 0.04 |
| (R)TELPPEK(L) | 100% | 28.6 | 18.6 | 1.82 | 407.2213 | 812.4280 | 2 | -0.0002 | -0.27 |
| (R)IATGQIAGVDK(D) | 100% | 31.1 | 21.8 | 2.52 | 536.8033 | 1071.5921 | 2 | -0.0006 | -0.57 |
| (R)GVGcLDFSK(A) | 100% | 33.4 | 19.1 | 0.96 | 491.7368 | 981.4591 | 2 | 0.0000 | -0.03 |
| (K)TTVEPTPGK(G) | 98% | 30.7 | 20.5 | | 465.2505 | 928.4864 | 2 | -0.0004 | -0.43 |
| (R)EIEVPDQYGTIR(A) | 99% | 20.0 | 22.2 | 1.09 | 710.3607 | 1418.7068 | 2 | 0.0024 | 1.68 |
| (R)WFVLDAETR(D) | 100% | 37.0 | 21.9 | 2.08 | 568.7901 | 1135.5657 | 2 | -0.0006 | -0.51 |
| (K)LSLPQNETVADTTLTK(A) | 100% | 31.8 | 21.9 | 3.10 | 865.9627 | 1729.9109 | 2 | 0.0008 | 0.46 |

**C**

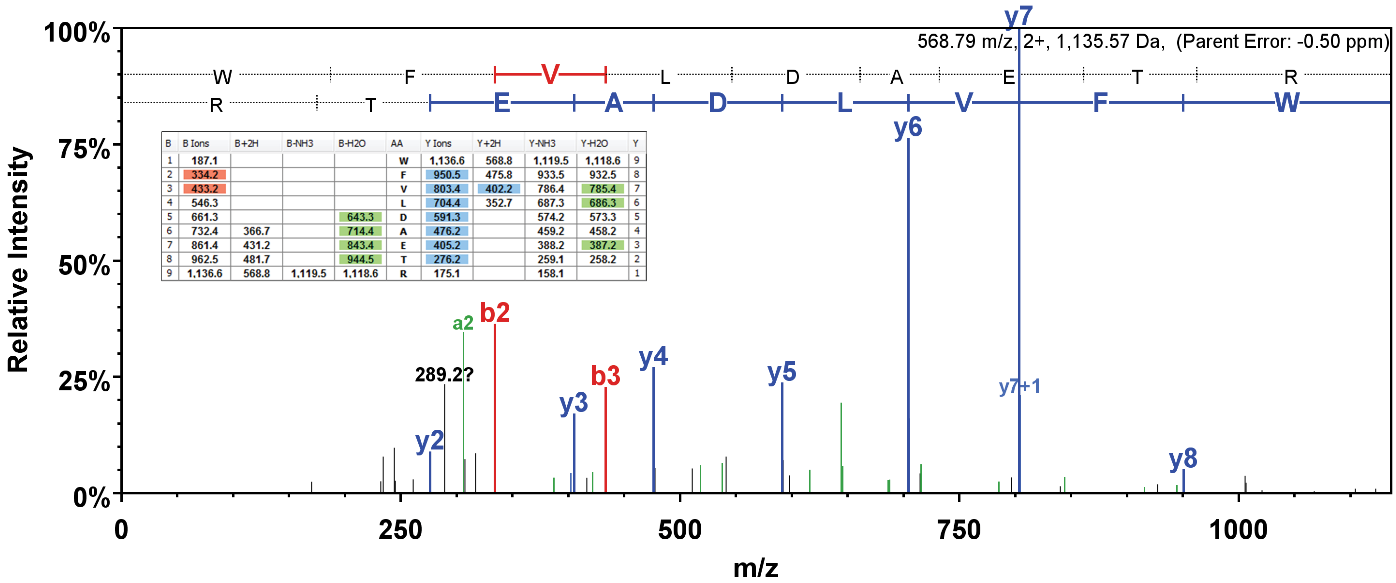


| B | B Ions | B+2H | B-NH3 | B-H2O | AA | Y Ions | Y+2H | Y-NH3 | Y-H2O | Y |
|---|---|---|---|---|---|---|---|---|---|---|
| 1 | 187.1 | | | | W | 1,136.6 | 568.8 | 1,119.5 | 1,118.6 | 9 |
| 2 | 334.2 | | | | F | 950.5 | 475.8 | 933.5 | 932.5 | 8 |
| 3 | 433.2 | | | | V | 803.4 | 402.2 | 786.4 | 785.4 | 7 |
| 4 | 546.3 | | | | L | 704.4 | 352.7 | 687.3 | 686.3 | 6 |
| 5 | 661.3 | | | 643.3 | D | 591.3 | | 574.2 | 573.3 | 5 |
| 6 | 732.4 | 366.7 | | 714.4 | A | 476.2 | | 459.2 | 458.2 | 4 |
| 7 | 861.4 | 431.2 | | 843.4 | E | 405.2 | | 388.2 | 387.2 | 3 |
| 8 | 962.5 | 481.7 | | 944.5 | T | 276.2 | | 259.1 | 258.2 | 2 |
| 9 | 1,136.6 | 568.8 | 1,119.5 | 1,118.6 | R | 175.1 | | 158.1 | | 1 |

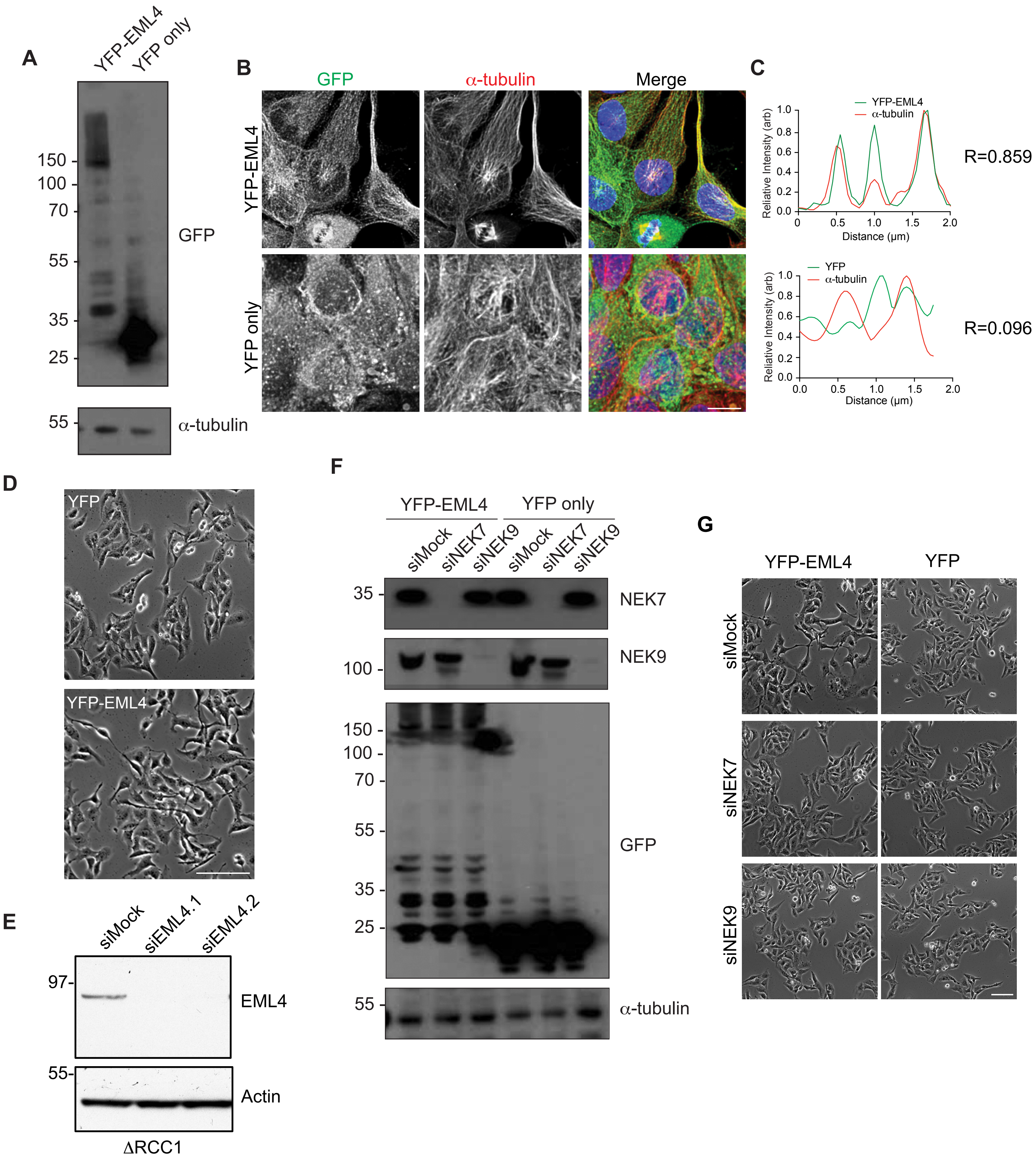
A
YFP-EML4
YFP only
150
100
70
55
35
25
GFP
55
α-tubulin
B
GFP
α-tubulin
Merge
YFP-EML4
YFP only
C
YFP-EML4
α-tubulin
Relative Intensity (arb)
Distance (µm)
R=0.859
YFP
α-tubulin
R=0.096
D
YFP
YFP-EML4
E
siMock
siEML4.1
siEML4.2
97-
EML4
55-
Actin
ΔRCC1
F
YFP-EML4
YFP only
siMock
siNEK7
siNEK9
siMock
siNEK7
siNEK9
35
NEK7
100
NEK9
150
100
70
55
35
25
GFP
55
α-tubulin
G
YFP-EML4
YFP
siMock
siNEK7
siNEK9

**A**

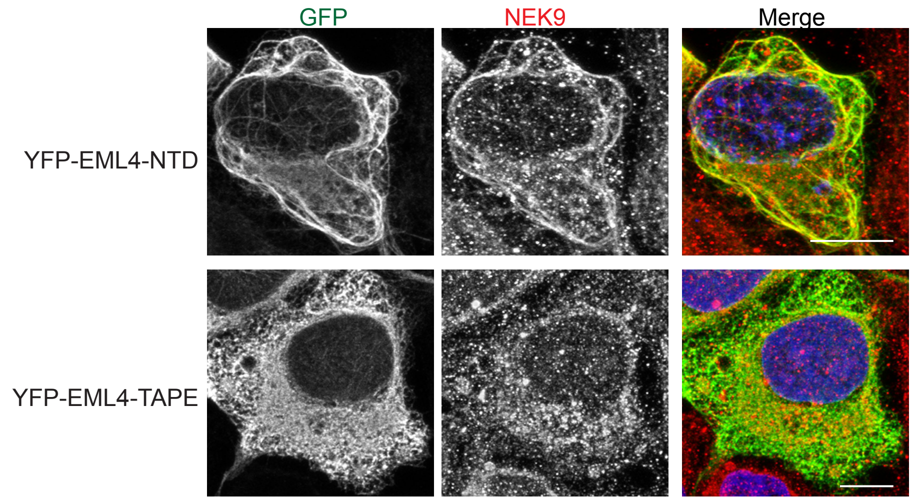
GFP
NEK9
Merge
YFP-EML4-NTD
YFP-EML4-TAPE

**B**

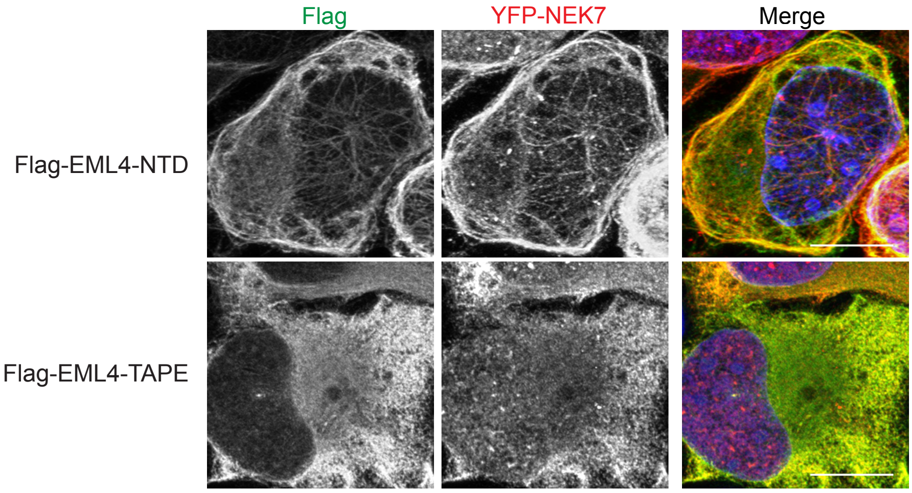
Flag
YFP-NEK7
Merge
Flag-EML4-NTD
Flag-EML4-TAPE

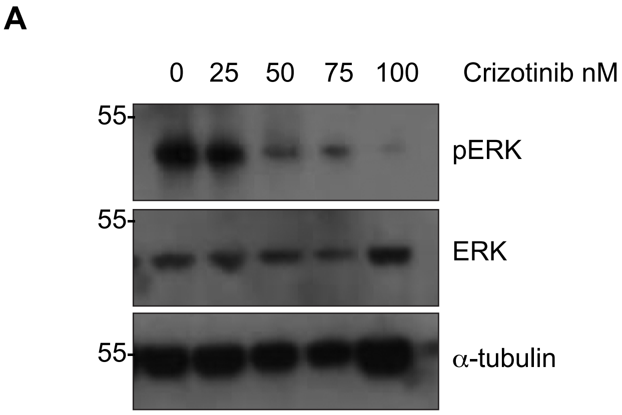
A
0
25
50
75
100
Crizotinib nM
55
pERK
55
ERK
55
α-tubulin

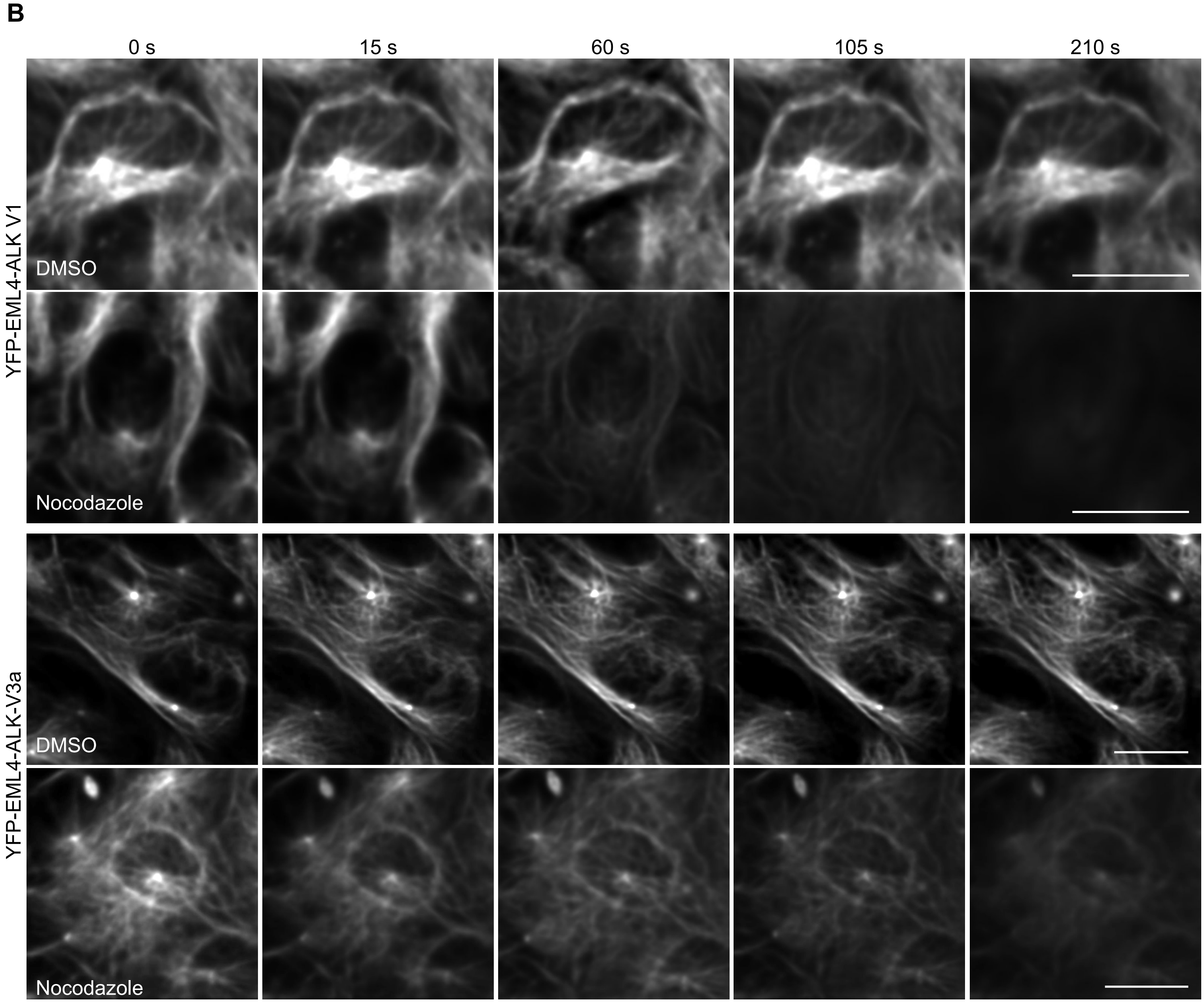
B
0 s
15 s
60 s
105 s
210 s
YFP-EML4-ALK V1
DMSO
Nocodazole
YFP-EML4-ALK-V3a
DMSO
Nocodazole

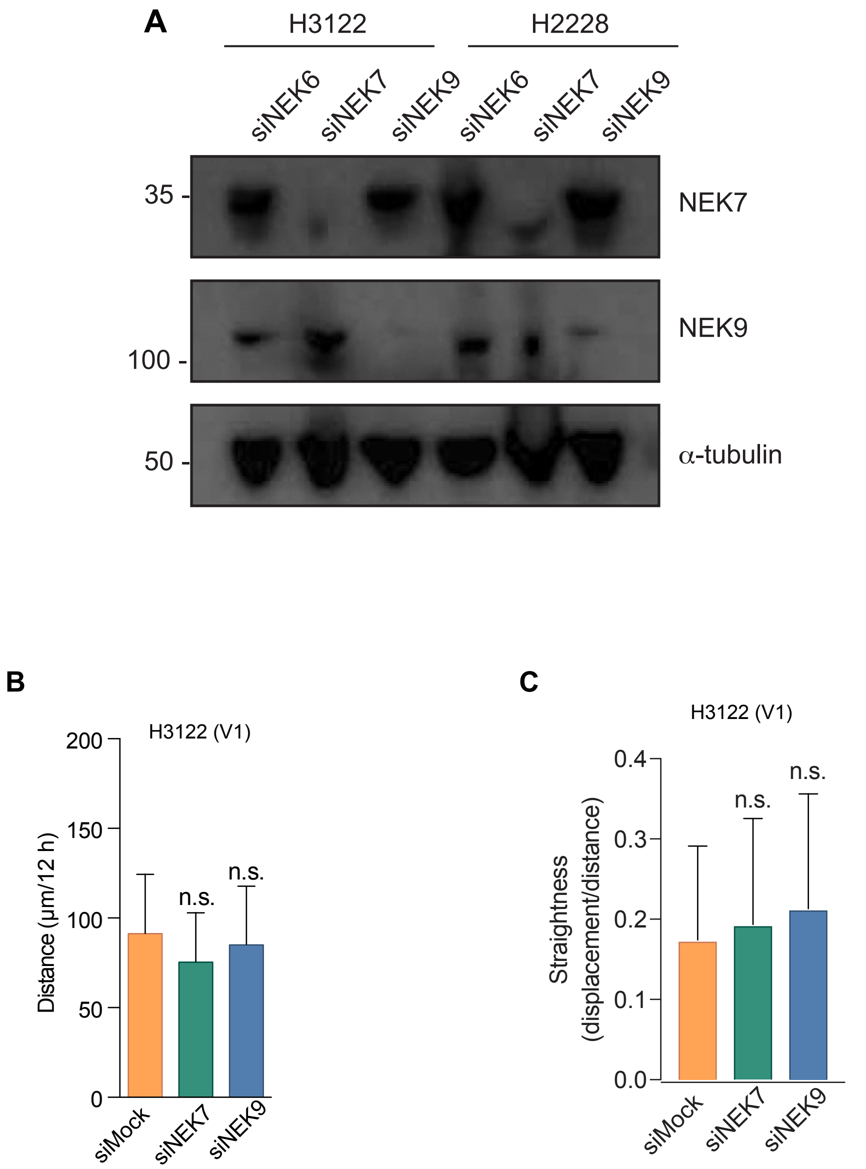
A
H3122
H2228
siNEK6
siNEK7
siNEK9
siNEK6
siNEK7
siNEK9
35 -
NEK7
NEK9
100 -
50 -
α-tubulin
B
H3122 (V1)
Distance (µm/12 h)
200
150
100
50
0
n.s.
n.s.
siMock
siNEK7
siNEK9
C
H3122 (V1)
Straightness (displacement/distance)
0.4
0.3
0.2
0.1
0.0
n.s.
n.s.
siMock
siNEK7
siNEK9

| Parameters | $N$ (%) |
|---|---|
| Sex | |
| Male | 27 (49.1) |
| Female | 28 (50.9) |
| Age in years, median (range) | 54 (27 - 79) |
| Smoking history | |
| Never | 37 (67.3) |
| ≤10 pack-years | 8 (14.5) |
| >10 pack-years | 10 (18.2) |
| Number of previous chemotherapy | |
| 0 | 38 (69.1) |
| 1 | 11 (20.0) |
| ≥ 2 | 6 (10.9) |
| Follow up in months, median (range) | 15 (1 - 54) |

| Parameters - *N* (%) | Score 1+ (*N*=22) | Score 2+/3+ (*N*=33) | *P* value* |
|---|---|---|---|
| Sex | | | 1.000 |
| Male | 11 (50) | 16 (48.5) | |
| Female | 11 (50) | 17 (51.5) | |
| Age in years, median (range) | 55 (27 - 75) | 54 (37 – 76) | 0.911 |
| Smoking history | | | 0.390 |
| Never | 14 (63.6) | 23 (69.7) | |
| ≤10 pack-years | 5 (22.7) | 3 (9.1) | |
| >10 pack-years | 3 (13.6) | 7 (21.2) | |
| Number of previous therapy | | | 0.071 |
| 0 | 13 (59.1) | 25 (75.8) | |
| 1 | 4 (18.2) | 7 (21.2) | |
| ≥ 2 | 5 (22.7) | 1 (3.0) | |
| Follow up in months, median (range) | 15 (2 - 50) | 15 (1 - 54) | 0.486 |

.
